# The motive cocktail in children’s altruistic behaviors

**DOI:** 10.64898/2026.03.18.712612

**Authors:** Xiaoyan Wu, Xiangjuan Ren, Jean-Claude Dreher, Chao Liu

## Abstract

Children frequently intervene in social conflicts by punishing violators or helping victims, yet the motivational mechanisms underlying such third-party altruistic behavior remain poorly understood. It remains unclear how children balance fairness concerns against self-interest, how these motivations interact with intervention costs and impact on outcomes, and whether gender and individual differences reflect distinct motivational structures. Here, we applied the motive cocktail model, which assumes that altruistic behavior arises from multiple prosocial motives, to dissociate motivations underlying third-party interventions. We studied 229 children aged 8–12 years (123 boys), an age when fairness and inequality aversion are reliably expressed. The third-party intervention task manipulated inequality between others, the personal cost of intervention, its impact on outcomes, and the form of intervention (punishment versus helping). Children intervened more as inequality increased and less as intervention costs rose, indicating a trade-off between moral benefits and self-interest. Gender differences emerged only under high-cost and high-impact conditions, with boys engaging in more punishment interventions. The motive cocktail model outperformed alternative models and revealed that boys showed stronger aversion to disadvantageous inequality and a greater tendency to reverse victims’ disadvantage than girls. Clustering analyses further identified distinct motivational profiles within each gender. These findings demonstrate that children’s third-party altruistic behavior is governed by multiple dissociable motives. This study provides a mechanistic account of how social motivations are organized and weighted during late childhood.

## Introduction

Children exhibit prosocial behaviors from an early age, including helping others, sharing resources, and responding to unfairness (1–3). Beyond direct interactions, children also intervene as third parties when observing unfair allocations between others, either by punishing violators (4, 5) or helping victims (6, 7). Such third-party interventions are thought to play a critical role in maintaining social norms and supporting cooperation in human societies (8–10). Children across cultures incur personal costs to address unfairness between others, suggesting a tendency to favor fairness (2, 11). Extending this developmental trajectory, evidence from infancy indicates that sensitivity to prosocial and norm-relevant actions emerges prior to language and explicit norm learning (12–14).

Research shows that children are highly sensitive to inequality and can distinguish between disadvantageous and advantageous outcomes for themselves (4, 15–17). As third-party observers, children increasingly intervene in response to unequal outcomes between others (2, 7, 18). These interventions can take multiple forms. Punishment may deter norm violations by reducing the violator’s payoff, whereas helping directly improves the welfare of the victim (6, 7). Both forms of intervention increase as inequality becomes more severe (2, 11). Children’s willingness to intervene is also constrained by the personal cost of acting (2, 5, 19). However, it remains unclear whether reduced intervention reflects aversion to personal costs or dissatisfaction with the degree of inequality reduction.

A further theoretical question concerns the motives underlying children’s altruistic decisions. Behaviors such as punishing a violator or helping a victim are often interpreted as expressions of fairness concerns. However, the same intervention may also reflect multiple underlying motivations, including restoring balance between social partners, reducing disadvantageous outcomes, or promoting efficiency (6, 7). Some accounts suggest that children’s altruistic behavior largely reflects the internalization or imitation of socially learned moral norms (20). In contrast, other work suggests that children’s prosocial actions arise from multiple cognitive and affective processes that jointly shape social decision-making (21, 22). Distinguishing among these possibilities requires experimental paradigms that allow different motivational components to be formally dissociated (23).

Another unresolved issue concerns gender and individual differences in children’s third-party interventions. Previous findings regarding gender differences in prosocial behavior have been mixed. Some studies report gender differences in costly punishment, with boys sometimes engaging in punishment more frequently than girls (2), whereas others find little evidence for stable gender differences across societies (11, 24). More generally, children show substantial heterogeneity in how they respond to social conflicts, with some favoring punishment and others preferring helping strategies (6, 7, 25). These observations raise the possibility that apparent gender differences may not reflect differences in overall altruism but rather differences in how underlying social motives are weighted when responding to inequality.

Here, we address these questions by combining a third-party intervention task with computational modeling in children aged 8–12 years, a developmental period during which fairness concerns and inequality aversion are reliably expressed in social decision-making (4, 15, 26). The task systematically manipulated the degree of inequality between others, the personal cost of intervention, the impact of intervention on outcomes, and the form of intervention (punishment versus helping). We first examined how these contextual factors influence children’s intervention decisions and whether these effects differ by gender. We then applied hierarchical Bayesian modeling to test whether children’s altruistic behavior is best explained by the *motive cocktail* framework, which assumes that altruistic behavior arises from the integration of multiple prosocial motives (9). Finally, clustering analyses were used to characterize individual differences in motivational profiles. This approach allowed us to examine how multiple motives jointly shape children’s altruistic behavior and how motivational weights vary across contexts, gender, and individuals.

## Results

### Children’s altruistic behaviors increased with others’ inequality and decreased with intervention cost

Consistent with previous findings, children’s altruistic behaviors increased with higher levels of inequality between the violator and the victim (*b* (regression coefficient from Generalized Linear Mixed Model) = 0.408, 95% confidence interval (CI) [0.251, 0.565], *p* < 0.001; pos hoc: 9:1 vs. 8:2 (*t*(228) = 3.10, *p*_corr_ = 0.004); 8:2 vs. 7:3 (*t*(228) = 0.61, *p*_corr_ = 0.542, Bayes factor (BF_10_) = 0.30); 7:3 vs. 6:4 *t*(228) = 5.19, *p*_corr_ < 0.001; 6:4 vs. 5:5: *t*(228) = 5.55, *p*_corr_ < 0.001; Fig. 1C).

**Fig. 1.**
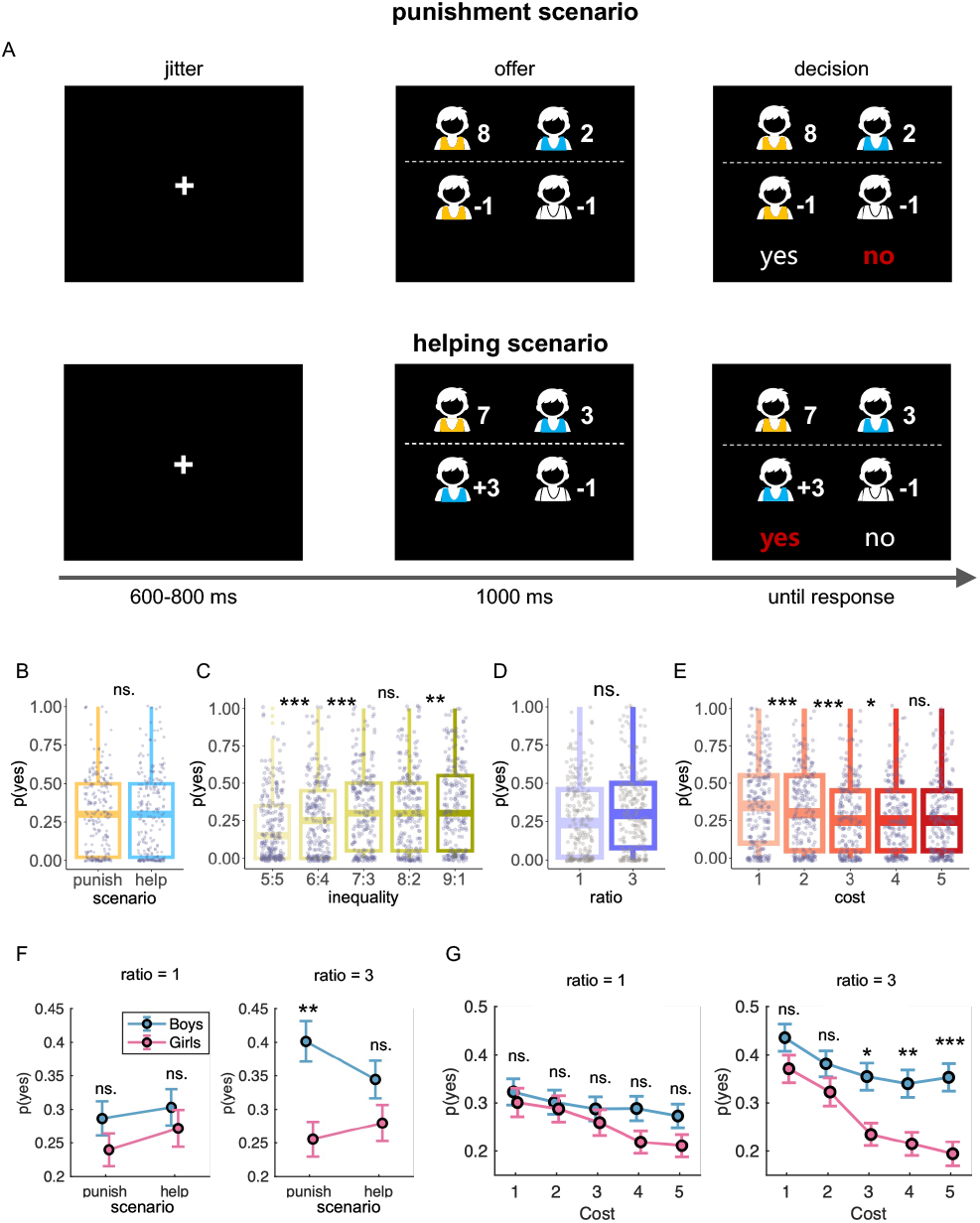
Third-party intervention task and behavioral results. (A) Schematic of the intervention task for punishment (top row) and helping (bottom row) scenarios. In each trial, participants observed the outcome of a resource allocation game in which a violator (orange shirt) allocated tokens between themselves and a victim (blue shirt), for example, 8 (to self):2 (to the victim). In the punishment scenario, participants (white shirt) could pay a cost (ranging from 1 to 5 tokens) to reduce the violator’s payoff (e.g., −1×cost or −3×cost tokens). In the helping scenario, participants could pay a cost to increase the victim’s payoff (e.g., +1×cost or +3×cost tokens). Participants then chose whether to accept (“yes”) or reject (“no”) the intervention offer. (B–E) Main effects of scenario (B), inequality (C), ratio (D), and cost (E) on intervention probability (i.e., the probability of choosing “yes”). Each gray dot represents one participant. Boxes indicate the 25th, 50th (median), and 75th percentiles; whiskers extend to 1.5× the interquartile range. (F) Interaction of scenario × ratio × gender. (G) Interaction of cost ratio gender. Each circle denotes group means across participants; error bars represent s.e.m. \*\*\**p* < 0.001; \*\**p* < 0.01; \**p* < 0.05, Holm–Bonferroni corrected; ns., not significant.

Children were also sensitive to the personal cost of intervening in others’ inequality. Higher personal costs reduced their probability of intervening (*b* = −0.465, 95% CI [−0.599, −0.331], *p* < 0.001; post hoc: 1 vs. 2: *t*(228) = 3.92, *p*_corr_ < 0.001; 2 vs. 3: *t*(228) = 4.19, *p*_corr_ < 0.001; 3 vs. 4: *t*(228) = 2.48, *p*_corr_ = 0.028; 4 vs. 5: *t*(228) = 1.07, *p*_corr_ = 0.285, BF_10_ = 0.31; Fig. 1E).

Children showed no preference for punishing the violator over helping the victim (*b* = −0.054, 95% CI [−0.422, 0.314], *p* = 0.773; Fig. 1B), nor for low-versus high-impact ratios (*b* = 0.112, 95% CI [−0.064, 0.288], *p* = 0.212; Fig. 1D). Age showed no significant effects (*b*_age_ = −0.223, 95% CI [−0.631, 0.185], *p* = 0.284; *b*_age× gender_ = 0.271, 95% CI [−0.242, 0.783], *p* = 0.301; see Supplementary Methods and Table S8). Given the narrow age range, age was not included in subsequent analyses.

### Boys exhibit more altruistic behavior than girls under high-impact ratio and high-cost conditions

The interaction of scenario × ratio× gender (*b* = 0.207, 95% CI [0.047, 0.366], *p* = 0.011) revealed that boys exhibited more punishment intervention than girls in the high-impact ratio condition (ratio = 3: *t*(227) = 3.60, 95% CI [0.066, 0.226], *p*_corr_ = 0.002). This gender effect was not observed under the low-impact ratio condition (ratio = 1: *t*(227) = 1.32, 95% CI [− 0.023, 0.117], *p*_corr_ = 0.377, BF_01_ = 1.14). In contrast, no reliable gender differences were found in help intervention at either ratio level (ratio = 3: *t*(227) = 1.66, 95% CI [−0.012, 0.142], *p*_corr_ = 0.297, BF_01_ = 0.73; ratio = 1: *t*(227) = 0.81, 95% CI [−0.045, 0.108], *p*_corr_ = 0.418, BF_01_ = 1.84; Fig. 1F).

The interaction of cost× ratio ×gender (*b* = 0.120, 95% CI [0.009, 0.230], *p* = 0.034) revealed that boys exhibited more intervention than girls only in the high-impact and high-cost conditions (cost = 3: *t*(227) = 3.20, 95% CI [0.046, 0.193], *p*_corr_ = 0.013; cost = 4: *t*(227) = 3.29, 95% CI [0.050, 0.199], *p*_corr_ = 0.010; cost = 5: *t*(227) = 4.13, 95% CI [0.083, 0.234], *p*_corr_ = 0.001). This gender difference was not observed in the low-cost conditions (cost = 1: *t*(227) = 1.61, 95% CI [− 0.015, 0.145], *p*_corr_ = 0.547, BF_01_ = 0.78; cost = 2: *t*(227) = 1.47, 95% CI [ −0.020, 0.137], *p*_corr_ = 0.568, BF_01_ = 0.94), nor under any cost level in the low-impact ratio conditions (cost = 1: *t*(227) = 0.54, 95% CI [− 0.057, 0.101], *p*_corr_ = 1, BF_01_ = 2.16; cost = 2: *t*(227) = 0.38, 95% CI [−0.059, 0.087], *p*_corr_ = 1, BF_01_ = 2.31; cost = 3: *t*(227) = 0.78, 95% CI [−0.043, 0.100], *p*_corr_ = 1, BF_01_ = 1.88; cost = 4: *t*(227) = 2.00, 95% CI [0.001, 0.138], *p*_corr_ = 0.325, BF_01_ = 0.41; cost = 5: *t*(227) = 1.81, 95% CI [−0.005, 0.129], *p*_corr_ = 0.430, BF_01_ = 0.57; Fig. 1G). No significant main effects were observed for scenario (*b* = −0.054, 95% CI [−0.422, 0.314], *p* = 0.773; Fig. 1B), gender (*b* = 0.541, 95% CI [−0.098, 1.180], *p* = 0.097) or ratio (*b* = 0.112, 95% CI [−0.064, 0.289], *p* = 0.212; Fig. 1D).

### The motive cocktail model reveals distinct motives underlying gender differences in children’s altruistic behavior

Children’s altruistic behaviors were assumed to be driven by motivations to consider self-interest (SI), reduce self-centered inequality aversion (SCI; controlled by *α* for disadvantageous inequality aversion and *β* for advantageous inequality aversion), to reduce victim-centered inequality aversion (VCI; *γ*), to increase efficiency concern (EC; *ω*), and to reverse inequality between the violator and the victim via reversal preference (RP; *κ*). We incorporated two inequality discounting components induced by intervention costs: *η*_keep_, capturing the discounting of inequality between others before intervention, and *η*_act_, capturing the discounting of residual inequality between others after intervention (see 2A for conceptual illustration). Parameters were estimated using a hierarchical Bayesian framework with Markov chain Monte Carlo (MCMC) sampling, with individual-level parameters assumed to be drawn from gender-specific group-level distributions (see Supplementary Methods for details).

Model construction followed our previous work (9), starting from the assumption that individuals only consider SI, and progressively adding distinct prosocial motives. We also constructed a baseline model with fixed intervention probabilities and two heuristic models. In total, ten models were compared (see Supplementary Methods). Pareto-smoothed importance sampling leave-one-out cross-validation (PSIS–LOO) showed that, consistent with our previous findings in adults, children’s altruistic behaviors were best explained by the motive cocktail model with seven distinct prosocial motives (Figs. 2B-D).

**Fig. 2.**
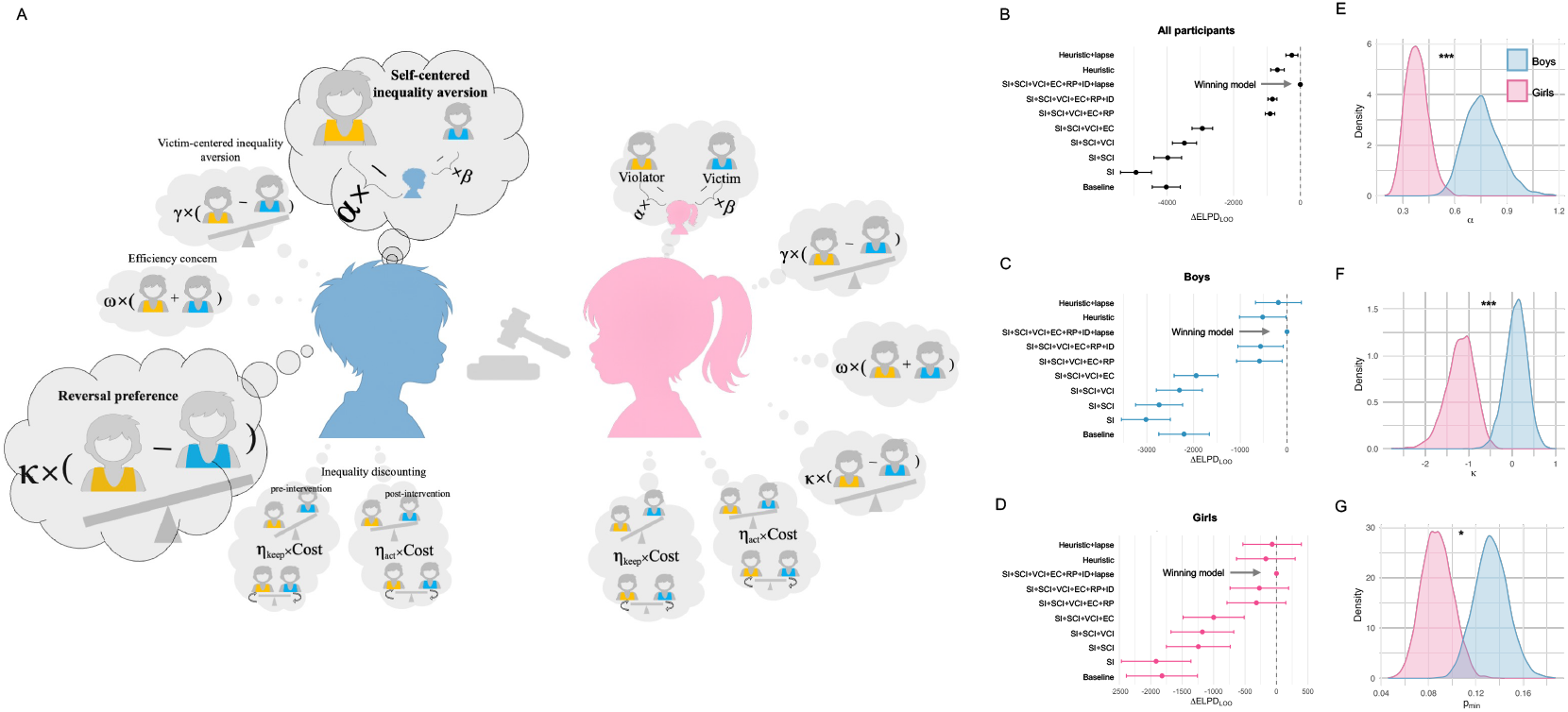
The motive cocktail model and gender differences in prosocial motives. (A) Conceptual illustration of the motive cocktail model. (B–D) Model comparison using PSIS–LOO cross-validation for all participants (B), boys (C), and girls (D). Points indicate differences in expected log predictive density (ΔELPD_LOO_) relative to the best-performing model (set to zero), with horizontal error bars showing *±*1 standard error. The dashed vertical line marks the winning model in each group. (E–G) Posterior distributions of group-level parameters for boys (blue) and girls (pink): disadvantageous inequality aversion (*α*) (E), reversal preference (*κ*) (F), and lower-bound intervention probability (*p*_min_) (G). Shaded areas indicate posterior densities. Asterisks denote the posterior probability of a gender difference (\**p* < 0.05; \*\*\**p* < 0.001, two-tailed).

Posterior parameter comparisons indicated that boys were more averse to disadvantageous inequality (*α*; mean difference = 0.38, 95% credible interval (CrI) [0.15, 0.62], *p* < 0.001; Fig. 2E) and showed a stronger preference for reversing inequality by placing the violator in a disadvantaged position (*κ*; mean difference = 1.29, 95% CrI [0.52, 2.12], *p* < 0.001; Fig. 2F) compared with girls. Boys also exhibited a higher baseline tendency to intervene across conditions (*p*_min_; mean difference = 0.05, 95% CrI [0.01, 0.08], *p* = 0.016; Fig. 2G; see Supplementary Fig. 2 for all parameters and Supplementary Fig. 5 for the robustness of gender patterns across age groups).

### Gender and individual differences in behavioral patterns

To characterize individual heterogeneity in altruistic behaviors, an unsupervised clustering analysis (K-means) was applied separately to boys and girls based on their intervention probabilities across 100 experimental conditions. Boys were best classified into two subgroups, whereas girls were best classified into three subgroups (Fig. 3A). *Distinct behavioral patterns among boys*. Boys in Cluster 1 (*n* = 37, 30%) showed a generally high probability of intervention across conditions, particularly under high inequality and high-impact ratio conditions (Fig. 3B). Boys in Cluster 2 (*n* = 86, 70%) displayed consistently low intervention probability, with weak sensitivity to inequality, intervention cost and impact ratio (Fig. 3C). This pattern is similar to the *Rational Moralists* identified in our previous study of adults (9), who seek to achieve an acceptable standard of morality at the lowest cost.

**Fig. 3.**
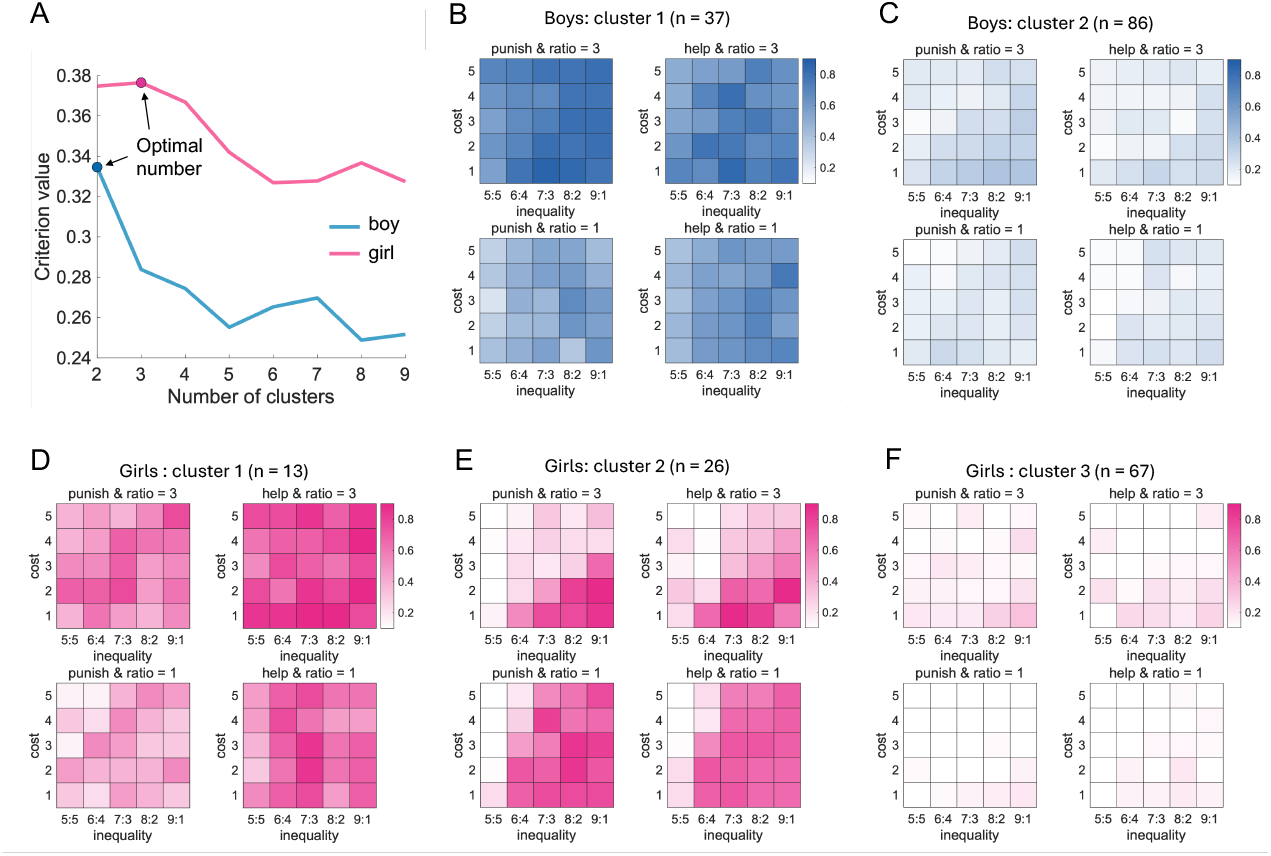
Gender and individual differences in behavioral patterns. (A) Silhouette coefficients across different numbers of clusters are shown to assess clustering quality. The silhouette coefficient quantifies the extent to which individuals are more similar to others within the same cluster than to those in different clusters, with higher values indicating stronger within-cluster consistency and better between-cluster separation. Based on this metric, the optimal number of clusters was two for boys and three for girls. (B–C) Behavioral patterns of the two boy clusters (blue), and (D–F) behavioral patterns of the three girl clusters (pink). Each panel depicts the average probability of choosing to intervene across experimental contexts for a given cluster. Within each panel, each cell represents a specific experimental condition. The x-axis indicates the degree of inequality between the violator and the victim (e.g., 8:2 indicates that the violator keeps 8 of 10 units and allocates 2 units to the victim), and the y-axis denotes the cost of intervention ranging from 1 to 5. Color intensity represents the probability of intervention, ranging from 0 to 1, where 0 indicates never intervening (i.e., choosing “no”) and 1 indicates always intervening (i.e., choosing “yes”) in that condition. In each panel, the left column corresponds to the punishment condition and the right column to the helping condition; the top row shows ratio = 3, and the bottom row shows ratio = 1. Blue and pink color scales represent boys and girls, respectively.

#### Greater heterogeneity in altruistic behavioral patterns among girls

Girls in Cluster 1 (*n* = 13, 12.26%) showed generally high intervention probabilities, especially in the helping scenario (Fig. 3D). The pattern is similar to the *Pragmatic Helpers* identified in our previous study of adults (9). Girls in Cluster 2 (*n* = 26, 24.52%) displayed a scenario-, inequality-, and cost-dependent pattern, with intervention probabilities increasing with inequality but declining sharply as intervention costs rose, particularly in the high-impact ratio condition (Fig. 3E). The pattern is similar to the *Justice Warriors* identified in our previous study of adults (9). Girls in Cluster 3 (*n* = 67, 63.20%) showed consistently low intervention probabilities, with weak sensitivity to inequality, intervention cost, and impact ratio (Fig. 3F). This pattern is similar to the *Rational Moralists* identified in our previous study of adults (9).

### Gender-specific motivational profiles underlying behavioral heterogeneity

We next compared cluster-specific profiles of the motive cocktail model parameters within each gender to elucidate the underlying internal mechanisms of the behavioral heterogeneity identified above.

Among boys, the two behavioral clusters identified in Figs. 3B–C corresponded to sharply distinct motivational profiles (Fig. 4A). Compared with Cluster 2, boys in Cluster 1 were less averse to disadvantageous inequality (*α*: Wilcoxon rank sum test, *r* = 0.279, *p* = 0.012), more averse to victim-centered inequality (*γ*: *r* = 0.566, *p* < 0.001), and showed a stronger reversal preference (*κ*: *r* = 0.559, *p* < 0.001). Boys in Cluster 1 also exhibited higher upper- and lower-bound intervention probabilities (*p*_max_: *r* = 0.376, *p* < 0.001; *p*_min_: *r* = 0.393, *p* < 0.001; Fig. 4C and Supplementary Table S5)

**Fig. 4.**
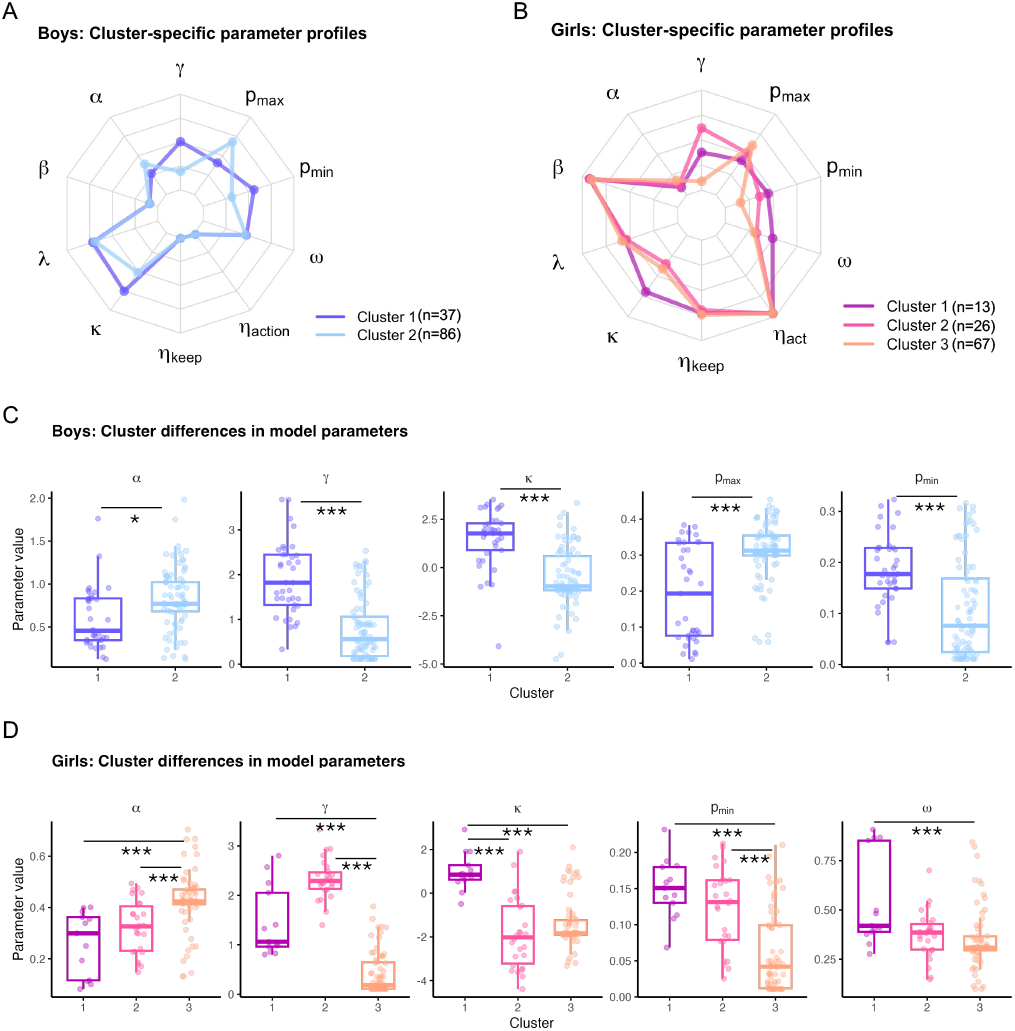
Cluster-specific parameter profiles in boys and girls. (A) Cluster-specific parameter profiles in boys and (B) girls. Radar plots illustrate cluster-specific profiles of model parameters after min–max normalization across all participants (i.e., each parameter was scaled to the [0, 1] range using the global minimum and maximum across all participants). Each axis corresponds to one model parameter, and values closer to the outer edge indicate relatively larger parameter magnitudes. Separate radar plots are shown for boys (top) and girls (bottom), with different colors representing distinct clusters identified within each gender (boys: two clusters; girls: three clusters). For each gender, boxplots show the distribution of subject-level posterior means for selected parameters across clusters. Individual dots represent single participants, boxes indicate the interquartile range, and the horizontal line denotes the median. Statistical comparisons were conducted within each gender. (C) For boys (two clusters), Wilcoxon rank-sum tests were used. (D) For girls (three clusters), Kruskal–Wallis tests were first applied, followed by post hoc pairwise comparisons. All reported post hoc *p* values were corrected for multiple comparisons using the Holm–Bonferroni procedure across parameters (\**p* < 0.05; \*\*\**p* < 0.001, two-tailed).

Among girls, the three behavioral clusters were associated with distinct motivational profiles (Fig. 4B). Cluster 1 (characterized as *Pragmatic Helpers*) was motivated by a strong preference for reversing inequality between the violator and the victim compared with both Cluster 2 (*κ*: post hoc Dunn test following Kruskal–Wallis, *r* = 0.489, *p* < 0.001) and Cluster 3 (*κ*: *r* = 0.427, *p* < 0.001). This pattern resembled the *Pragmatic Helpers* identified in our previous adult study (9). Girls in Cluster 1 were also more motivated by efficiency concern than Cluster 3 (*ω*: *r* = 0.343, *p* < 0.001) and exhibited generally higher tendency to intervene compared with Cluster 3 (*p*_min_: *r* = 0.439, *p* < 0.001). Cluster 2 (characterized as *Justice Warriors*) was motivated to reduce the inequality suffered by the victim more than girls in Cluster 3 (*γ*: *r* = 0.766, *p* < 0.001), and exhibited a generally high baseline intervention tendency (*p*_min_: *r* = 0.423, *p* < 0.001). Cluster 3 (characterized as *Rational Moralists*) exhibited an overall low tendency to intervene. They were motivated to reduce disadvantageous inequality for themselves compared with both Cluster 1 (*α*: *r* = −0.425, *p* < 0.001) and Cluster 2 (*α*: *r* = −0.364, *p* < 0.001), but barely cared about the inequality suffered by the victim compared with both Cluster 1 (*γ*: *r* = 0.396, *p* < 0.001) and Cluster 2 (*γ*: *r* = 0.766, *p* < 0.001; Fig. 4D and Supplementary Table S6 and Table S7).

## Discussion

Children’s altruistic behaviors were strongly increased with the degree of inequality between others and decreased with the personal cost of intervening, indicating that children are balancing moral benefits against self-interest when responding to social inequality. This pattern is consistent with evidence that children across cultures are willing to incur costs to address unfairness (11, 27), and that third-party altruistic behaviors reflect a trade-off between inequality aversion and self-interest (28). Relatedly, recent work shows that adolescents also act against self-interest in social dilemmas (29), suggesting that prosocial decisions balance moral motives and personal costs. Children in general showed no overall preference for punishing the violator over helping the victim, and no overall gender differences across conditions. Only at high impact ratios (i.e., when intervention is worthwhile) did boys prefer punishment over girls. Under conditions of high intervention costs and a high impact ratio (i.e., when intervention is powerful), boys also showed more third-party intervention. These results suggest that altruistic behavior is guided by multiple considerations and varies across individuals, even at early developmental stages (1). Such findings argue against interpreting gender differences in terms of “higher levels of altruism or empathic concern” and instead support a context-triggered account. The results are consistent with evidence that gender differences in prosocial behavior are not universal across societies (24), but emerge selectively under high-risk or high-stakes conditions (30).

As we found in our previous study in adults (9), the *Motive Cocktail Model*, which combines seven distinct prosocial motives, also provided the best account of children’s altruistic behaviors, suggesting that a multi-dimensional motivational architecture is already present in childhood. Children’s altruistic behaviors were jointly motivated by self-interest considerations, reductions in self-centered inequality, reductions in victim-centered inequality (in which sensitivity to the victim’s inequality is discounted by personal intervention costs), increased efficiency concerns, and inequality reversal between the violator and the victim. These findings challenge the view that children’s altruistic behavior primarily reflects the imitation of adult moral norms (20). Instead, these findings support the idea that children possess a complex and dissociable set of social motives early in development, consistent with evidence that multiple cognitive and affective processes jointly shape children’s prosocial behavior (21, 22).

Computational modeling further specified the internal motivational sources of gender differences in children’s altruistic behavior. Compared with girls, boys were more motivated to reduce disadvantageous inequality for themselves, showed a stronger reversal preference, and exhibited a higher baseline intervention probability, a motivational profile consistent with their greater tendency to engage in punitive intervention under high-conflict conditions (31, 32). This pattern is broadly consistent with previous findings showing that boys tend to respond more aggressively to conflict than girls (33). Our computational framework reveals distinct motivational profiles among boys.

Two distinct motivational profiles were further identified among boys. Subgroup 1 showed a high tendency to intervene (*p*_min_), and was more strongly motivated to reduce inequality between others (*γ*) and to reverse the relative advantage between violators and victims (*κ*) by third-party intervention. In contrast, Subgroup 2, the majority of boys, were primarily driven by the reduction of disadvantageous inequality for themselves (*α*), rather than by concerns about inequality between others. These findings indicate that boys’ punitive intervention in social conflicts reflects a clear differentiation between intervention motives and how self-versus other-oriented inequality concerns are weighted.

Girls exhibited greater heterogeneity in their motivational structure, clustering into at least three strategy types that closely resembled prosocial intervention profiles previously described in adults. However, these similarities should not be interpreted as evidence for more mature altruistic motivational development in girls, as our previous adult study has not directly examined gender differences. Subgroup 1 showed a high tendency to intervene (*p*_min_) and was primarily motivated to reverse the relative advantage between violators and victims (*κ*), reflecting a pragmatic orientation toward restoring social balance through third-party intervention. Subgroup 2 was characterized by a strong motivation to reduce inequality suffered by the victim (*γ*), accompanied by a generally high baseline intervention tendency. Subgroup 3 exhibited an overall low tendency to intervene, with interventions mainly driven by the reduction of disadvantageous inequality for themselves, and relatively weak concern for inequality experienced by others. Together, these findings indicate that girls’ third-party intervention in social conflicts is supported by multiple dissociable motivational profiles, reflecting differentiated emphases on status reversal, victim-oriented fairness, and self-oriented inequality concerns (34, 35).

By combining behavioral analyses with computational modeling, we demonstrate that children’s third-party punishment and helping behaviors are governed by multiple distinct social motives rather than a single equality-driven process. Gender differences in altruistic behavior are context-dependent and become visible under high-impact and high-cost conditions. This reflects gender differences in the weighting of different altruistic motivations. Children’s strategies exhibit similarities to altruistic types previously identified in adults, suggesting developmental continuity in social motivation. The task in this study is easy for children to understand and enables computational modeling of children’s prosocial decisions beyond what has been possible in much of the previous literature.

In sum, the present study demonstrates that children’s third-party altruistic intervention is not driven by a single moral impulse, but combines multiple prosocial motives. These motives are structurally organized within individuals during childhood and account for context-dependent gender differences as well as substantial individual heterogeneity in altruistic strategies. By introducing the motive cocktail framework to developmental research, our findings provide a mechanistic account of children’s moral decision-making and highlight both the continuity and flexibility of social motivation across development. More broadly, this work illustrates the value of computational approaches for uncovering the processes underlying complex social behaviors early in life.

## Materials and Methods

A total of 229 children participated in this experiment. All participants were primary school students from Grades 3 to 6 in Beijing, aged between 8 and 12 years (*M* ± *SD* = 9.74 ± 0.96; 123 boys). Participants’ ages were calculated in months and converted into years by dividing the number of months by 12 and rounding to two decimal places (e.g., 10 years and 6 months was recorded as 10.5 years). Sample size was based on our prior work using similar designs and practical considerations, and was not determined by an a priori power analysis.

The study was approved by the Ethics Committee of Beijing Normal University (ethics approval number: IRB A 0003 2020001) and was conducted in accordance with established ethical standards for research involving human participants, including the principles outlined in the Declaration of Helsinki. Prior to participation, written informed consent was obtained from children’s parents or legal guardians, as well as from the children, and participation was entirely voluntary.

### Third-party intervention task

We adapted the “third-party intervention task” used in adult altruistic behavior research by Wu et al. (2024) for children by simplifying the task structure. In the task, children played as a third party in two scenarios: punishing a violator and helping a victim. In each scenario, the inequality between the violator and the victim (5:5 [violator’s payoff: victim’s payoff], 6:4, 7:3, 8:2, and 9:1) and the intervention cost (1 to 5) were manipulated across five levels. The impact ratio had two levels (1 and 3). Both allocation and intervention offers were presented at the same time to reduce the cognitive load on children. Stimuli were presented until the children made a choice.

### Experimental procedure

The experiment was conducted in a group setting, with children participating as class groups in the school computer lab. One experimenter provided an explanation of the task instructions, followed by the children independently reading the instructions and completing a comprehension test. Several experimenters were available to assist the children in understanding the task and to answer any questions regarding the rules. All children were required to pass the comprehension test before proceeding to practice trials. The comprehension test consisted of scenario-based calculation questions from both the punishment and helping conditions and required children to compute the final token amounts for all parties based on the initial allocation, intervention cost, and impact ratio. Answers had to be generated as fill-in-the-blank calculations rather than selected from multiple-choice options. Children proceeded only after answering all questions correctly. Nearly all children provided correct answers, indicating a clear understanding of the task rules.

The experimental task consisted of two blocks: one involving the reduction of the violator’s payoff (punishment scenario) and the other involving an increase in the victim’s payoff (helping scenario). The block order was randomized across participants. Each block contained 50 trials, with the trial order being fully randomized. A 30-second break was given between the two blocks. The total task duration was approximately 20 minutes. Children were informed that 10% of the trials would be randomly selected, and the tokens they retained in those trials would be exchanged for stationery gifts. Different token amounts corresponded to different rewards, including pencils, pens, pencil cases, and stationery sets.

### Behavioral analysis. Generalized Linear Mixed Model (GLMM)

We applied a generalized linear mixed model (GLMM) to analyze the behavioral choices of the 229 children. In the model, the dependent variable was the participant’s choice (1 for “yes”, 0 for “no”), which followed a Bernoulli distribution. The independent variables included scenario (categorical: punishment or helping), degree of unfairness (continuous: violator’s money minus victim’s money), intervention cost (continuous: levels 1 to 5), impact ratio (continuous: levels 1 and 3), and trial number (continuous: trials 1 to 100). The model included random effects for each participant, consisting of a random intercept and random slopes for each independent variable.

In the model analyzing reaction times, the dependent variable was reaction time (in seconds), and the independent variables were the same as those included in the GLMM. In both models, random effects additionally included participants’ gender and age. All continuous variables were standardized prior to analysis, and the reported values correspond to standardized *β* coefficients. Significant effects and interactions were further examined using post hoc tests, with *p* values corrected for multiple comparisons using the Bonferroni adjustment.

### Linear Mixed Model (LMM)

We also conducted linear mixed-effects modeling on participants’ reaction times (in seconds), assuming a normal distribution. The independent variables in this model were identical to those used in the GLMM.

### Clustering Analysis

To identify distinct behavioral patterns, we performed a clustering analysis on children’s intervention decisions. K-means clustering was applied to the behavioral data, using Hamming distance as the similarity metric to account for the binary nature of the responses. Clustering quality was evaluated using the silhouette coefficient. For this analysis, each participant’s behavior was represented as a 100-dimensional binary vector, corresponding to the 100 experimental conditions in the third-party intervention and bystander paradigm, with each condition coded as “yes” (1) or “no” (0).

### Modeling

#### Motive cocktail model

We modeled intervention decisions using the *motive cocktail* model, which integrates multiple motivations within a unified utility-based framework. On each trial *t*, subject *s* chose whether to intervene (*y*_*st*_ = 1) or not (*y*_*st*_ = 0). Choices were assumed to be driven by comparisons between the subjective utilities of intervention (choosing “yes”) and non-intervention (choosing “no”), with an explicit lapse component to account for random responding.

#### Payoff structure

On trial 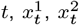, and 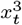 represent the payoffs to the violator, victim, and third-party (i.e., the participant), respectively. Intervention incurred a cost *c*_*t*_ to the third-party and affected payoffs according to the scenario-specific rule with impact ratio *r*_*t*_.

Payoffs under intervention were given by:

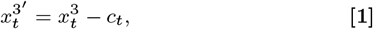

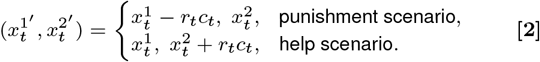

Under non-intervention, payoffs remained unchanged.

#### Self-interest (SI)

SI was defined as the third party’s own payoff before intervention (SI) and after intervention (SI’):

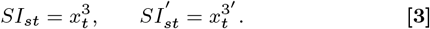

#### Self-centered inequality (SCI)

SCI assumes that children’s altruistic behavior is motivated by aversion to payoff inequality between the self and others. Accordingly, SCI quantifies payoff inequality between the self (i.e., the participant) and the two other parties, including disadvantageous inequality (DI; the self having less than others) and advantageous inequality (AI; the self having more than others):

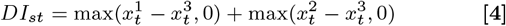

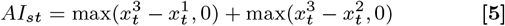

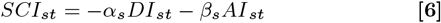

where *α*_*s*_ and *β*_*s*_ denote subject-specific aversion to disadvantageous and advantageous inequality, respectively.

Accordingly, when the third party chooses “yes”, DI, AI, and SCI change as a function of 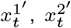, and 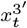, denoted 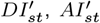, and 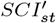, respectively.

#### Victim-centered inequality (VCI)

VCI assumes that children’s altruistic behavior is driven by aversion to payoff inequality between others. Accordingly, VCI quantifies payoff inequality between the violator and the victim:

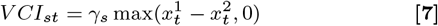

where *γ*_*s*_ denotes subject-specific aversion to violator-centered inequity. Accordingly, when the third-party chooses to “yes”, VCI changes as a function of 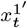 and 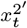, denoted 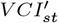.

#### Efficiency concern (EC)

EC assumes that children’s altruistic behavior is driven by increasing others’ payoffs and quantifies the total payoff of others (the violator and the victim):

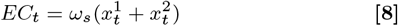

where *ω*_*s*_ denotes the subject-specific weight placed on increases in the total payoff of others. Accordingly, when the third-party chooses to “yes”, EC changes as a function of 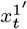 and 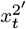, denoted 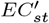.

#### Reversal preference (RP)

RP assumes that children’s altruistic behavior is driven by reversing the payoff advantage of the violator, and quantifies situations in which the victim receives a higher payoff than the violator after third-party intervention:

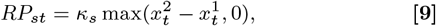

where *κ*_*s*_ denotes the subject-specific weight placed on reversal preference. When the third-party chooses to “yes”, RP changes as a function of 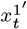 and 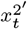, denoted 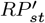.

#### Inequality discounting (ID)

ID assumes that sensitivity to VCI is systematically discounted by the intervention cost. Accordingly, two cost-discounted terms were defined for the non-intervention and intervention options, respectively:

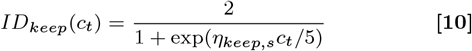

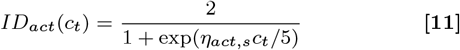

where *η*_keep,*s*_ and *η*_act,*s*_ denote subject-specific parameters governing the degree of cost-dependent discounting of VCI under non-intervention and intervention, respectively. As the intervention cost increases, sensitivity to VCI decreases.

#### Utility functions

Combining all components, the subject utility for the non-intervention option (i.e., choosing “no”), 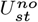, is defined as:

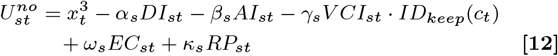

And the subject utility for the intervention option (i.e., choosing “yes”), 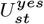, is defined as:

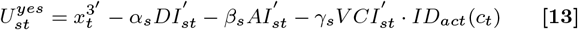

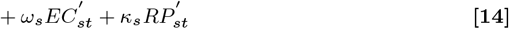

#### Choice rule with lapse

Utility-based choice probability is given by:

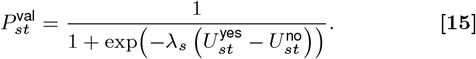

where *λ*_*s*_ denotes the subject-specific inverse temperature parameter, with larger values indicating more deterministic choices.

Final choice probabilities incorporated subject-specific lapse bounds:

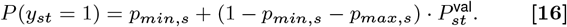

Here, *p*_*min,s*_ captures a baseline tendency to intervene independent of utility, whereas *p*_*max,s*_ captures a baseline tendency to refrain from intervention. The remaining probability mass governs value-based decision-making.

#### Hierarchical bayesian inference

See Supplementary Methods for details.

#### Model comparison and validation

See Supplementary Methods, Supplementary Fig. 3 and Fig. 4 for details.

## Author Contributions

Conceptualization: X.W. and C.L. Investigation, Data curation, Formal analysis, Software, and Visualization: X.W. Methodology: X.W., X.R., J.-C.D., and C.L. Writing-original draft: X.W. Writing-review & editing: X.W., X.R., J.-C.D., and C.L. Funding acquisition and Supervision: C.L.

## Data, Materials, and Software Availability

All data, analysis code, computational modeling scripts, and task materials are available at https://github.com/xiaoyanwu2024/child-altruism.git.

## ACKNOWLEDGMENTS

We thank all participants and experimenters for their assistance with data collection.

