## Supporting Information for "The motive cocktail in children’s altruistic behaviors"

#### Section 1: Supplementary Methods

**Age effects on altruistic behaviors.** To examine potential age effects, we fitted a generalized linear mixed-effects model (GLMM) predicting altruistic actions. Fixed effects included block, inequality, cost, ratio, gender, age, and the age × gender interaction. All predictors were standardized (z-scores) prior to analysis. The model included participant-level random intercepts and random slopes for block, inequality, cost, ratio, age, and gender, and was estimated with a binomial distribution and logit link. Full results are reported in Table S8.

**Model construction.** To investigate the motivations underlying intervention decisions, we formalized models based on different assumptions. Following our previous work (1), we started with a baseline model (Model 1). Models 2–7 assumed that decisions were driven by different combinations of altruistic motives. Model 2 included only self-interest, and Models 3–7 each included one additional altruistic motive. We kept a motive only if it improved model fit according to Pareto-smoothed importance sampling leave-one-out cross-validation (PSIS-LOO). Model fit continued to improve until all motives were included in the motive cocktail model. Based on this model, we further introduced a lapse parameter to allow bounded choice probabilities, resulting in Model 8. In addition, we constructed two heuristic models (Models 9 and 10) for comparison.

Across all models, decisions were modeled at the trial level within a hierarchical Bayesian framework. Individual-level parameters were assigned prior distributions and drawn from gender-specific group-level distributions. We define the notation used in the model below. Let $s = 1, \dots, S$ index subjects and $t = 1, \dots, T$ index trials. On each trial $t$, subject $s$ chose whether to intervene ($y_{st} = 1$) or not ($y_{st} = 0$).

On trial $t$, the violator, victim, and the third-party (i.e., the participant) received payoffs denoted by $x_t^1$, $x_t^2$, and $x_t^3$, respectively. The value of $x_t^3$ was 5 in all conditions. Choosing to intervene incurred a cost $c_t$ to the third-party and affected either the payoff of the violator (under the punishment scenario) or that of the victim (under the help scenario) according to an impact ratio $r_t$. If the third-party decided to intervene, the payoffs were transformed as:

$$x_t^{3'} = x_t^3 - c_t \tag{1}$$

$$(x_t^{1'}, x_t^{2'}) = \begin{cases} (x_t^1 - r_t c_t,\ x_t^2), & \text{punishment scenario} \\ (x_t^1,\ x_t^2 + r_t c_t), & \text{help scenario} \end{cases} \tag{2}$$

In non-intervention trials, payoffs remained unchanged.

***Model 1: Baseline.*** Model 1 served as a lower-bound benchmark and assumed no sensitivity to trial-level variables. Each subject was characterized by a constant probability of intervention across trials:

$$P(y_{st} = 1) = p_s \tag{3}$$

where $p_s \in (0, 1)$ captured the subject-specific baseline tendency to intervene.

For Models 2–8, we specified a subjective utility for intervening ($U_{st}^{\text{yes}}$) and not intervening ($U_{st}^{\text{no}}$).

***Model 2: Self-interest (SI).*** Model 2 assumed that decisions were driven solely by self-interest. The utility of each option depended only on the participant's own payoff:

$$U_{st}^{\text{yes}} = x_t^{3'}, \quad U_{st}^{\text{no}} = x_t^3. \tag{4}$$

Choices were generated via a logistic response model, which was introduced in Model 2 and shared by Models 3–8:

$$P(y_{st} = 1) = \frac{1}{1 + \exp(-\lambda_s (U_{st}^{\text{yes}} - U_{st}^{\text{no}}))} \tag{5}$$

where $\lambda_s > 0$ denotes subject-specific decision sensitivity. Observed choices were modeled as:

$$y_{st} \sim \text{Bernoulli}(P(y_{st} = 1)) \tag{6}$$

***Model 3: SI + Self-Centered Inequity (SCI).*** Model 3 extended the SI model by adding SCI relative to both the violator and the victim. SCI was defined with respect to the difference between others' payoffs and the third-party's payoff. Disadvantageous inequality aversion (DI) captured situations in which others earned more than the third-party:

$$\mathrm{DI}_{st} = \max(x_t^1 - x_t^3, 0) + \max(x_t^2 - x_t^3, 0) \quad [7]$$

whereas advantageous inequality (AI) captured situations in which the third-party earned more than others:

$$\mathrm{AI}_{st} = \max(x_t^3 - x_t^1, 0) + \max(x_t^3 - x_t^2, 0) \quad [8]$$

The utility of intervening for subject $s$ on trial $t$ was defined as:

$$U_{st}^{\text{no}} = x_t^3 - \alpha_s\, \mathrm{DI}_{st} - \beta_s\, \mathrm{AI}_{st} \quad [9]$$

where $\alpha_s$ and $\beta_s$ denote subject-specific aversion to disadvantageous and advantageous inequality, respectively.

Utility under **intervention** was defined analogously using untransformed payoffs. Disadvantageous and advantageous inequality were computed as:

$$\mathrm{DI}_{st}^{'} = \max(x_t^{1'} - x_t^{3'}, 0) + \max(x_t^{2'} - x_t^{3'}, 0) \quad [10]$$

$$\mathrm{AI}_{st}^{'} = \max(x_t^{3'} - x_t^{1'}, 0) + \max(x_t^{3'} - x_t^{2'}, 0) \quad [11]$$

Accordingly, the utility of non-intervention was:

$$U_{st}^{\text{yes}} = x_t^{3'} - \alpha_s\, \mathrm{DI}_{st}^{'} - \beta_s\, \mathrm{AI}_{st}^{'} \quad [12]$$

Choice probabilities were generated according to the response model introduced in Model 2.

***Model 4: SI + SCI + Victim-Centered Inequity (VCI).*** Model 4 further added VCI, which captures inequity between the violator and the victim. In the third-party intervention task, the victim's payoff was always less than or equal to that of the violator. Accordingly, VCI was defined as the payoff advantage of the violator over the victim:

$$\mathrm{VCI}_{st} = \max\big(x_t^1 - x_t^2,\, 0\big) \quad [13]$$

where $x_t^1$ and $x_t^2$ denote the violator's and victim's payoffs before intervention.

For the **intervention** option, VCI was defined as VCI$'$:

$$\mathrm{VCI}_{st}^{'} = \max\Big(x_t^{1'} - x_t^{2'},\, 0\Big) \quad [14]$$

Accordingly, the utility functions were given by:

$$U_{st}^{\text{no}} = x_t^3 - \alpha_s \mathrm{DI}_{st} - \beta_s \mathrm{AI}_{st} - \gamma_s \mathrm{VCI}_{st} \quad [15]$$

$$U_{st}^{\text{yes}} = x_t^{3'} - \alpha_s \mathrm{DI}_{st}^{'} - \beta_s \mathrm{AI}_{st}^{'} - \gamma_s \mathrm{VCI}_{st}^{'} \quad [16]$$

where $\gamma_s$ denotes subject-specific aversion to violator-centered inequity. Choice probabilities were generated according to the response model introduced in Model 2.

***Model 5: SI + SCI + VCI + Efficiency Concern (EC).*** Model 5 extended Model 4 by adding an EC motive, defined as increases in the total payoff of others (the violator and the victim):

$$\mathrm{EP}_t = x_t^1 + x_t^2 \quad [17]$$

$$\mathrm{EP}_t^{'} = x_t^{1'} + x_t^{2'} \quad [18]$$

Accordingly, the utility functions were given by:

$$U_{st}^{\text{no}} = x_t^3 - \alpha_s \mathrm{DI}_{st} - \beta_s \mathrm{AI}_{st} - \gamma_s \mathrm{VCI}_{st} + \omega_s \mathrm{EP}_{st} \quad [19]$$

$$U_{st}^{\text{yes}} = x_t^{3'} - \alpha_s \mathrm{DI}_{st}^{'} - \beta_s \mathrm{AI}_{st}^{'} - \gamma_s \mathrm{VCI}_{st}^{'} + \omega_s \mathrm{EP}_{st}^{'} \quad [20]$$

where $\omega_s$ denotes the subject-specific weight placed on increases in the total payoff of others. Choices were generated using the logistic response model introduced in Model 2.

***Model 6: SI + SCI + VCI + EC + Reversal Preference (RP).*** Model 6 extended Model 5 by adding a RP, defined as situations in which the victim received a higher payoff than the violator after third-party's intervention. Reversal preference was defined as the positive payoff difference between the victim and the violator:

$$\mathrm{RP}_t = \max\left(x_t^2 - x_t^1,\, 0\right) \tag{21}$$

Under non-intervention, $\mathrm{RP}_t$ was always zero and thus had no effect on utility; it is written here for notational symmetry.

$$\mathrm{RP}_t^{'} = \max\left(x_t^{2'} - x_t^{1'},\, 0\right) \tag{22}$$

Accordingly, the utility functions were given by:

$$U_{st}^{\text{no}} = x_t^3 - \alpha_s \mathrm{DI}_{st} - \beta_s \mathrm{AI}_{st} - \gamma_s \mathrm{VCI}_{st} + \omega_s \mathrm{EC}_{st} + \kappa_s \mathrm{RP}_{st} \tag{23}$$

$$U_{st}^{\text{yes}} = x_t^{3'} - \alpha_s \mathrm{DI}_{st}^{'} - \beta_s \mathrm{AI}_{st}^{'} - \gamma_s \mathrm{VCI}_{st}^{'} + \omega_s \mathrm{EC}_{st}^{'} + \kappa_s \mathrm{RP}_{st}^{'} \tag{24}$$

where $\kappa_s$ denotes the subject-specific weight placed on reversal preference. Choices were generated using the logistic response model introduced in Model 2.

***Model 7: SI + SCI + VCI + EC + RP + Inequality Discounting (ID).*** Model 7 extended Model 6 by allowing VCI to be discounted as a function of intervention cost. Two cost-dependent discounting functions were defined for the non-intervention and intervention options, respectively:

$$\mathrm{ID}_{\text{keep}}(c_t) = \frac{2}{1 + \exp(\eta_{\text{keep},s} c_t / 5)} \tag{25}$$

$$\mathrm{ID}_{\text{act}}(c_t) = \frac{2}{1 + \exp(\eta_{\text{act},s} c_t / 5)} \tag{26}$$

Accordingly, the utility functions were given by:

$$U_{st}^{\text{no}} = x_t^3 - \alpha_s \mathrm{DI}_{st} - \beta_s \mathrm{AI}_{st} - \gamma_s \mathrm{VCI}_{st} \cdot \mathrm{ID}_{\text{keep}}(c_t) + \omega_s \mathrm{EC}_{st} + \kappa_s \mathrm{RP}_{st} \tag{27}$$

$$U_{st}^{\text{yes}} = x_t^{3'} - \alpha_s \mathrm{DI}_{st}^{'} - \beta_s \mathrm{AI}_{st}^{'} - \gamma_s \mathrm{VCI}_{st}^{'} \cdot \mathrm{ID}_{\text{act}}(c_t) + \omega_s \mathrm{EC}_{st}^{'} + \kappa_s \mathrm{RP}_{st}^{'} \tag{28}$$

where $\eta_{\text{keep},s}$ and $\eta_{\text{act},s}$ denote subject-specific parameters governing the degree of cost-dependent discounting of VCI under non-intervention and intervention, respectively. Choices were generated using the logistic response model introduced in Model 2.

***Model 8: SI + SCI + VCI + EC + RP + ID + Lapse.*** Model 8 extended Model 7 by adding subject-specific lapse parameters to allow bounded choice probabilities. The value-based choice probability for subject $s$ on trial $t$ was defined as:

$$P_{st}^{\text{val}} = \frac{1}{1 + \exp(-\lambda_s\,(U_{st}^{\text{yes}} - U_{st}^{\text{no}}))} \tag{29}$$

where $U_{st}^{\text{yes}}$ and $U_{st}^{\text{no}}$ were defined as in Model 7.

The final probability of choosing the intervention option was given by:

$$P(y_{st} = 1) = p_{min,s} + (1 - p_{min,s} - p_{max,s}) \cdot P_{st}^{\text{val}} \tag{30}$$

where, $p_{min,s}$ and $p_{max,s}$ denote subject-specific lower and upper bounds on the probability of choosing the intervention option, respectively. Choices were generated using the logistic response model introduced in Model 2.

***Model 9: Heuristic Model.*** Model 9 specified a heuristic decision rule in which intervention choices were modeled as a function of salient trial-level features, without explicit utility decomposition. For subject $s$ on trial $t$, the probability of choosing the intervention option was given by:

$$P(y_{st}=1)=\frac{1}{1+\exp(-\,(\theta_{s,0}+\theta_{s,1}\,c_t+\theta_{s,2}\,max(x_t^1-x_t^2,0)+\theta_{s,3}\,r_t+\theta_{s,4}\,s_t))} \quad [31]$$

where $\theta_{s,0}$ denotes a subject-specific intercept and $\theta_{s,k}$ $(k=1,\ldots,4)$ denote subject-specific feature weights.

The included features were intervention cost $(c_t)$, payoff difference between violator and victim $(x_t^1-x_t^2)$, impact ratio $(r_t)$, and scenario $(s_t)$ was coded as a binary indicator $(s_t=0$ for help, $s_t=1$ for punishment).

***Model 10: Heuristic feature-based model with lapse.*** Model 10 extended Model 9 by adding subject-specific lapse parameters to allow bounded choice probabilities. The heuristic choice probability for subject $s$ on trial $t$ was defined as:

$$z_{st}=\theta_{s,0}+\theta_{s,1}\,c_t+\theta_{s,2}\,max(x_t^1-x_t^2,0)+\theta_{s,3}\,r_t+\theta_{s,4}\,s_t \quad [32]$$

The probability of choosing the intervention option was then given by:

$$P(y_{st}=1)=p_{min,s}+(1-p_{min,s}-p_{max,s})\cdot\frac{1}{1+\exp(-z_{st})} \quad [33]$$

where $p_{min,s}$ and $p_{max,s}$ denote subject-specific lapse bounds.

**Hierarchical Bayesian estimation.** All models were estimated within a hierarchical Bayesian framework. Subject-level parameters were allowed to vary across individuals, with hierarchical priors defined at the group level by gender. Let $g(s)\in\{1,2\}$ denote the gender group of subject $s$ (female or male).

***Subject-level parameters.*** Depending on the model, each subject $s$ was characterized by a subset of the following parameters. $\alpha_s$, $\beta_s$, and $\gamma_s$ denote subject-specific aversion to self-centered disadvantageous inequality, self-centered advantageous inequality, and victim-centered inequity, respectively. $\omega_s$ denotes the subject-specific weight on increases in others' total payoff, and $\kappa_s$ denotes the subject-specific weight on inequality reversals between the violator and the victim. $\eta_{s,\text{keep}}$ and $\eta_{s,\text{act}}$ denote subject-specific parameters governing cost-dependent discounting of victim-centered inequity under the non-intervention and intervention options, respectively. $\lambda_s$ denotes the subject-specific decision sensitivity parameter governing choice stochasticity. Finally, $p_{\min,s}$ and $p_{\max,s}$ denote subject-specific lower and upper bounds on choice probabilities, respectively.

In heuristic Models 9 and 10, choices were instead governed by subject-specific feature weights $\theta_{s,k}$, which were estimated within the same hierarchical framework.

Parameters representing motivate weights $(\alpha_s,\beta_s,\gamma_s,\omega_s,\eta_{keep,s},\eta_{act,s})$ were constrained to be non-negative, consistent with their theoretical interpretation. The reversal preference parameter $\kappa_s$ was allowed to take both positive and negative values. Lapse parameters $(p_{\min,s}$ and $p_{\max,s})$ were constrained to the interval $[0,0.5]$.

***Group-level distributions.*** Each subject-level parameter $\theta_s$ was assumed to follow a normal distribution with a gender-specific group-level mean and standard deviation:

$$\theta_s\sim\mathcal{N}\big(\mu_{\theta,g(s)},\sigma_{\theta,g(s)}\big)\,. \quad [34]$$

where $g(s)$ denotes the gender group of subject $s$. Parameter constraints were enforced via bounds on the support of the corresponding parameters, as specified in the model implementation.

Group-level means and standard deviations were estimated separately for each gender. Normal priors were assigned to the group-level means, with prior means informed by parameter estimates from our previous study. Group-level standard deviations were assigned uniform priors bounded at zero, with upper limits set to a broad range informed by earlier estimates to allow sufficient flexibility in parameter estimation.

$$\begin{aligned}
\mu_{\alpha,g}&\sim\mathcal{N}(5,3), & \sigma_{\alpha,g}&\sim\mathcal{U}(0,6)\\
\mu_{\beta,g}&\sim\mathcal{N}(5,3), & \sigma_{\beta,g}&\sim\mathcal{U}(0,6),\\
\mu_{\gamma,g}&\sim\mathcal{N}(5,3), & \sigma_{\gamma,g}&\sim\mathcal{U}(0,6),\\
\mu_{\omega,g}&\sim\mathcal{N}(5,3), & \sigma_{\omega,g}&\sim\mathcal{U}(0,6),\\
\mu_{\kappa,g}&\sim\mathcal{N}(0,5), & \sigma_{\kappa,g}&\sim\mathcal{U}(0,6),\\
\mu_{\eta_{\text{keep}},g}&\sim\mathcal{N}(5,3), & \sigma_{\eta_{\text{keep}},g}&\sim\mathcal{U}(0,15),\\
\mu_{\eta_{\text{act}},g}&\sim\mathcal{N}(5,3), & \sigma_{\eta_{\text{act}},g}&\sim\mathcal{U}(0,15),\\
\mu_{\lambda,g}&\sim\mathcal{N}(10,5), & \sigma_{\lambda,g}&\sim\mathcal{U}(0,6),\\
\mu_{p_{\min},g}&\sim\mathcal{N}(0.25,0.1), & \sigma_{p_{\min},g}&\sim\mathcal{U}(0,0.5),\\
\mu_{p_{\max},g}&\sim\mathcal{N}(0.25,0.1), & \sigma_{p_{\max},g}&\sim\mathcal{U}(0,0.5).
\end{aligned}$$

All group-level means and standard deviations were assigned normal priors. Positivity and interval constraints were enforced through parameter bounds as specified in the model implementation.

***Estimation.*** Posterior inference was performed using Hamiltonian Monte Carlo sampling implemented in Stan via the `rstan` package in R. For each model, 4 Markov chains were run for 4,000 iterations, including 2000 warm-up iterations. Sampling used an adapt_delta of 0.95 and a maximum tree depth of 12. Model comparison was conducted using leave-one-out cross-validation based on subject-level log-likelihoods. Convergence diagnostics indicated satisfactory convergence for all reported parameters, with $\hat{R}$ values close to 1 and generally below the conservative threshold of 1.01 (Fig. S1).

**Model prediction.** To compare the predictive performance of the candidate models, we generated trial-level behavioral predictions for each model using posterior parameter estimates and evaluated their correspondence with observed behavior across experimental conditions. We considered ten models (m01–m10), spanning from simple baseline formulations to more structured models incorporating inequality aversion, efficiency concerns, role preferences, attention modulation, and lapse processes.

For each model, posterior samples were obtained from hierarchical Bayesian fits implemented in Stan. Subject-level posterior mean estimates were extracted for all free parameters (e.g., choice sensitivity, inequality aversion, efficiency weights, role preference coefficients, attention parameters, and lapse rates, depending on the model). Using these posterior means, we reconstructed each model's likelihood function and computed trial-wise predicted probabilities of intervention given the actual experimental inputs (violator and victim allocations, intervention cost, ratio, and block type).

To generate discrete model-predicted choices, we performed Bernoulli sampling at the trial level, drawing a binary intervention decision from a Bernoulli distribution parameterized by the predicted probability for each trial. These simulated choices were then aggregated across trials and subjects to obtain model-predicted mean intervention probabilities for each experimental condition. In parallel, empirical mean intervention probabilities were computed from the observed data by averaging intervention choices across all participants within the same conditions.

Model predictions and empirical data were compared across a total of 100 experimental conditions, defined by the combination of allocation unfairness levels, intervention costs, ratios, and punishment versus help contexts. The correspondence between observed and model-predicted mean intervention probabilities is shown in Supplementary Fig. S3.

### Section 2: Supplementary Results

This section reports additional statistical analyses that support the main findings. In particular, analyses of decision time were conducted to rule out potential speed–accuracy trade-offs underlying the observed behavioral effects.

**Model-based replication of behavioral effects.** To assess whether the hierarchical Bayesian model could account for the main behavioral findings, we generated trial-level predicted choices from the best-fitting model. Predicted choices were obtained by sampling from a Bernoulli distribution using the model-implied choice probability. The same statistical analyses applied to the behavioral data were then performed on the model-generated choices.

The model reproduced the main effect of inequality observed in the behavioral data. Predicted intervention rates increased as inequality became more extreme, with reliable differences between adjacent inequality levels (9:1 vs. 8:2: $t(228) = 3.00$, $p_{\text{Corr}} = 0.003$; 8:2 vs. 7:3: $t(228) = 3.71$, $p_{\text{Corr}} = 0.001$; 7:3 vs. 6:4: $t(228) = 6.95$, $p_{\text{Corr}} < 0.001$; 6:4 vs. 5:5: $t(228) = 4.09$, $p_{\text{Corr}} < 0.001$). The model also captured the main effect of intervention cost, showing a systematic decrease in predicted intervention probability as costs increased (cost 1 vs. 2: $t(228) = 5.59$, $p_{\text{Corr}} < 0.001$; cost 2 vs. 3: $t(228) = 3.61$, $p_{\text{Corr}} = 0.001$; cost 3 vs. 4: $t(228) = 3.52$, $p_{\text{Corr}} = 0.001$; cost 4 vs. 5: $t(228) = 2.55$, $p_{\text{Corr}} = 0.011$).

The model further reproduced the interaction between scenario, ratio, and gender. In the punishment scenario, a gender difference emerged under the high-ratio condition (ratio = 3), with boys showing a higher predicted likelihood of punishment than girls ($t(227) = 3.81$, 95% CI [0.072, 0.226], $p_{\text{Corr}} = 0.001$). No gender difference was observed under the low-ratio condition (ratio = 1; $t(227) = 1.35$, $p_{\text{Corr}} = 0.359$, $BF_{01} = 1.10$). In the helping scenario, predicted gender differences were not reliable at either ratio level (ratio = 1: $t(227) = 0.68$, $p_{\text{Corr}} = 0.497$, $BF_{01} = 2.01$; ratio = 3: $t(227) = 2.28$, $p_{\text{Corr}} = 0.071$, $BF_{01} = 0.24$).

A similar pattern was observed for the interaction between cost, ratio, and gender. Under the high-ratio condition (ratio = 3), boys showed higher predicted punishment rates than girls at medium to high cost levels (cost 3: $t(227) = 2.93$, $p_{\text{Corr}} = 0.026$; cost 4: $t(227) = 3.52$, $p_{\text{Corr}} = 0.005$; cost 5: $t(227) = 3.33$, $p_{\text{Corr}} = 0.008$). No reliable gender differences were found at lower cost levels or under any cost level in the low-ratio condition (all $p_{\text{Corr}} \geq 0.268$, $BF_{01} \geq 0.46$).

Overall, the model-generated choices recapitulated the main effects and interaction patterns observed in the behavioral data, while preserving conditions in which no reliable differences were present (see Supplementary Fig. S4).

### Section 3: Supplementary Figures

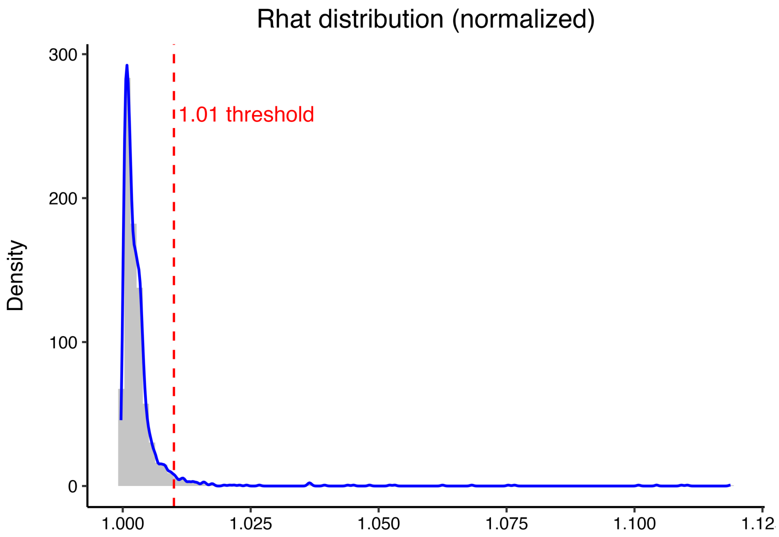


**Supplementary Figure 1. Convergence diagnostics for the best-fitting hierarchical Bayesian model (the motive cocktail model).** Normalized distribution of $\hat{R}$ values for all reported parameters. The vast majority of $\hat{R}$ values are clustered very close to 1.00 and remain below the conservative convergence threshold of 1.01 (red dashed line), indicating that the reported parameters achieved satisfactory convergence and were well mixed across MCMC chains.

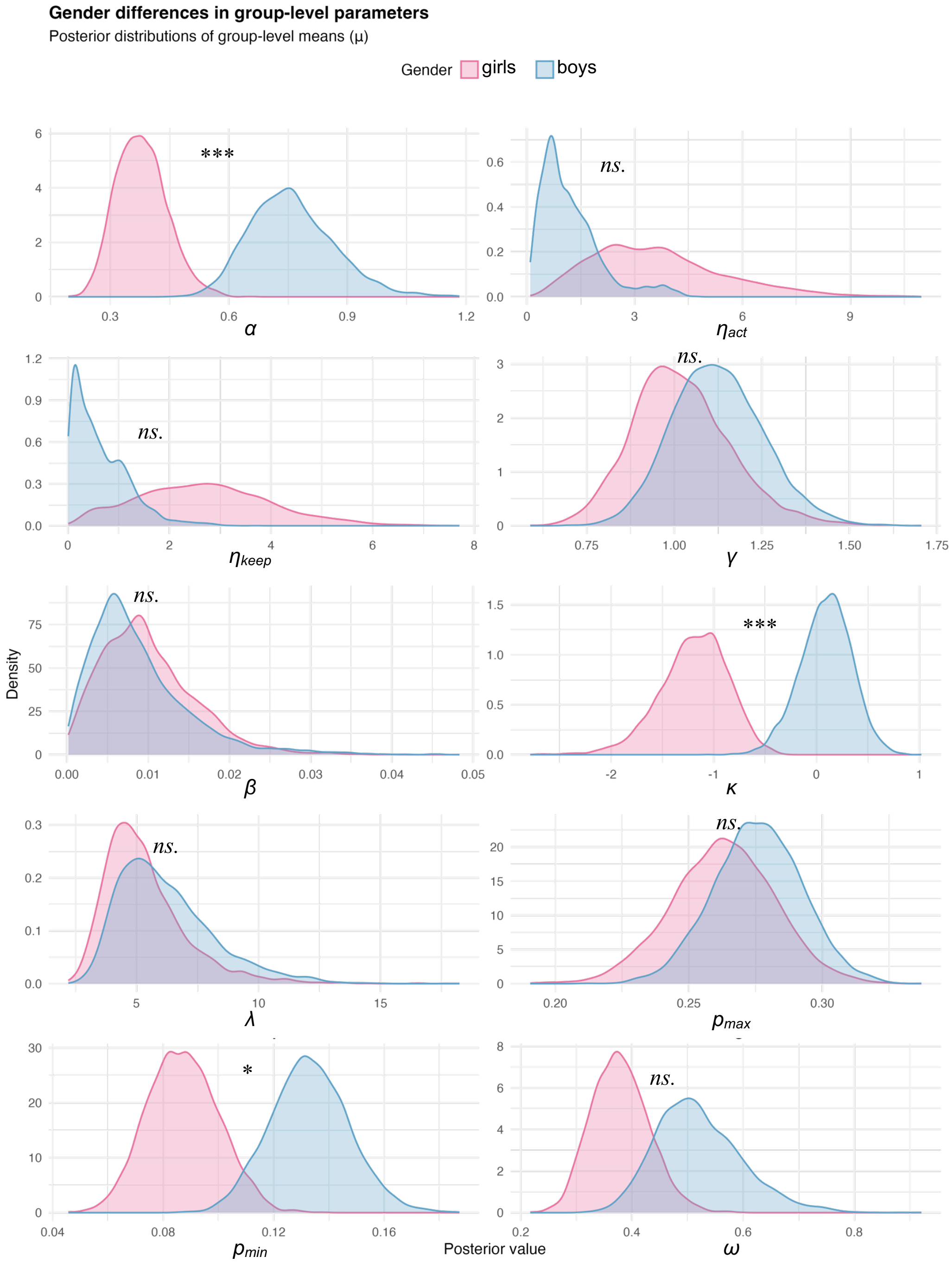


**Supplementary Figure 2. Gender differences in group-level parameters of the motive cocktail model.** Posterior distributions of group-level means for the ten parameters of the motive cocktail model, shown separately for boys (blue) and girls (red). Distributions are based on posterior samples from the hierarchical Bayesian model. Gender differences were assessed by computing the posterior distribution of the difference (boys minus girls) for each parameter. Significance markers indicate two-tailed posterior probabilities derived from these difference distributions (***$p < 0.001$, **$p < 0.01$, *$p < 0.05$; ns, not significant). Parameter symbols correspond to the model definitions: inequality sensitivity ($\alpha$), guilt sensitivity ($\beta$), inequity aversion ($\gamma$), choice sensitivity ($\lambda$), reciprocity preference ($\kappa$), cost modulation parameters ($\eta_k$, $\eta_a$), outcome utility weight ($\omega$), and lapse-rate bounds ($p_{\min}$, $p_{\max}$).

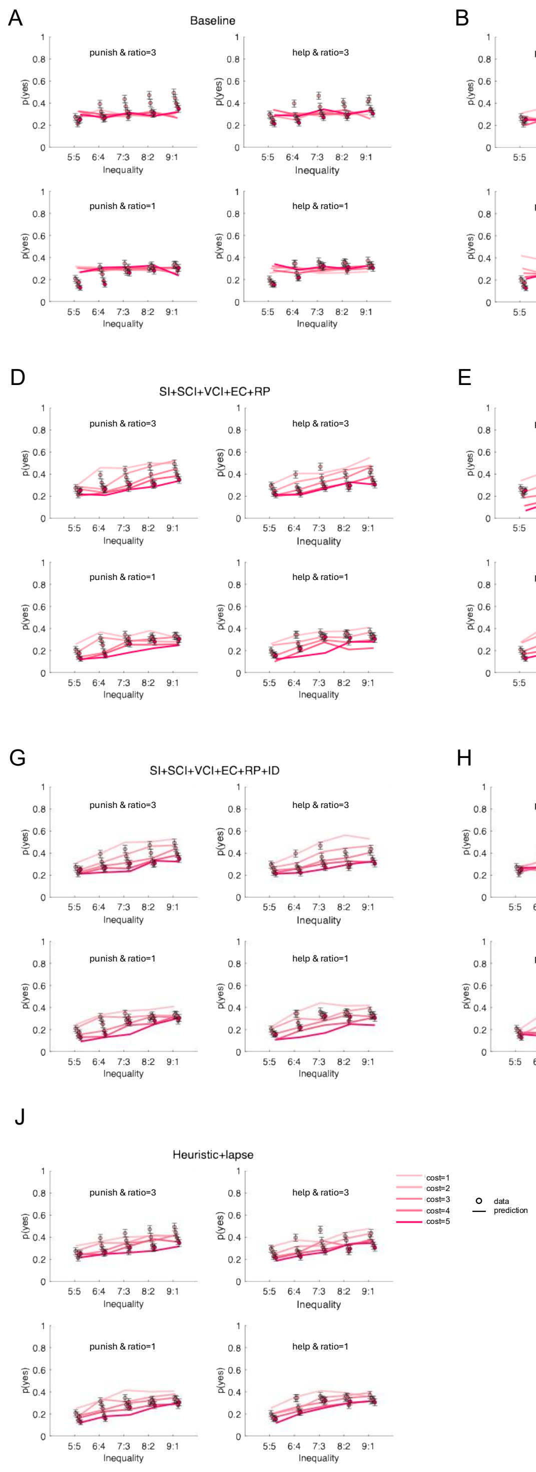


**Supplementary Figure 3. Observed and model-predicted intervention behavior across experimental conditions.** Panels A–J correspond to models m01–m10, respectively. The x-axis represents the degree of unfairness in the allocation between the violator and the victim, defined as the difference in their endowments. The y-axis shows the probability of choosing to intervene. Dots indicate the observed mean probability of intervention, with error bars representing $\pm 1$ standard error of the mean (SEM). Lines show the model-predicted mean probability of intervention based on posterior parameter estimates. Different colors denote the five levels of intervention cost. Within each panel, subplots are arranged by experimental condition: the left column shows the punishment condition and the right column shows the help condition; the top row corresponds to ratio = 3 and the bottom row corresponds to ratio = 1. This layout enables a direct comparison between observed behavior and model predictions across levels of unfairness, intervention cost, and task context.

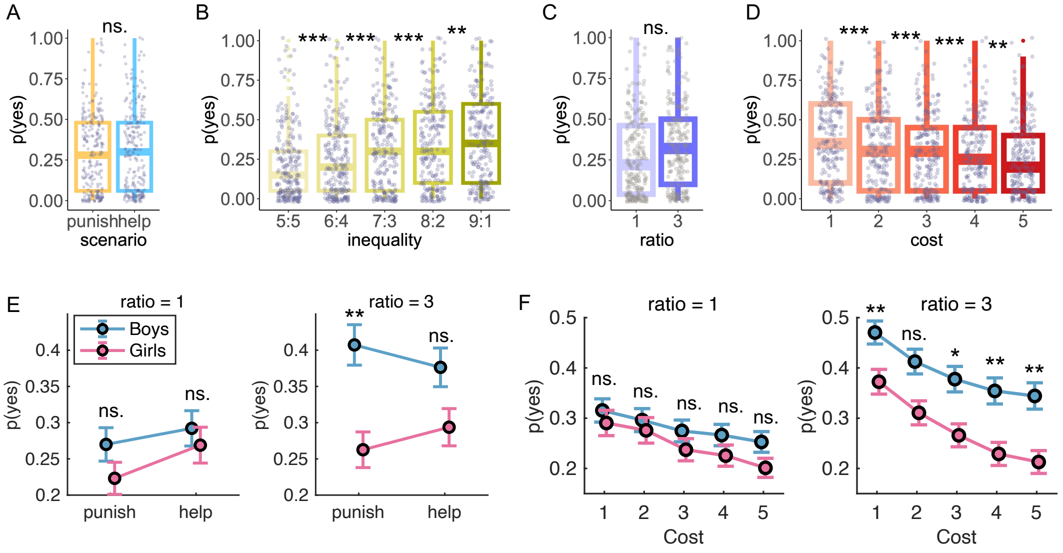


**Supplementary Figure 4. Reproduction of behavioral effects by the motive cocktail model.** Using posterior mean estimates of subject-level parameters from the hierarchical Bayesian generative model (the motive cocktail model), simulated behavioral data were generated for each participant to assess whether the model reproduced the main behavioral effects. Main effects of scenario (A), inequality (B), ratio (C), and cost (D) on intervention probability (i.e., the probability of choosing "yes"). Each gray dot represents one participant. Boxes indicate the 25th, 50th (median), and 75th percentiles; whiskers extend to 1.5 times the interquartile range. (E) Interaction of scenario $\times$ ratio $\times$ gender. (F) Interaction of cost $\times$ ratio $\times$ gender. Circles denote group means across participants; error bars represent s.e.m. ***$p < 0.001$; **$p < 0.01$; *$p < 0.05$ (Holm–Bonferroni corrected); ns, not significant.

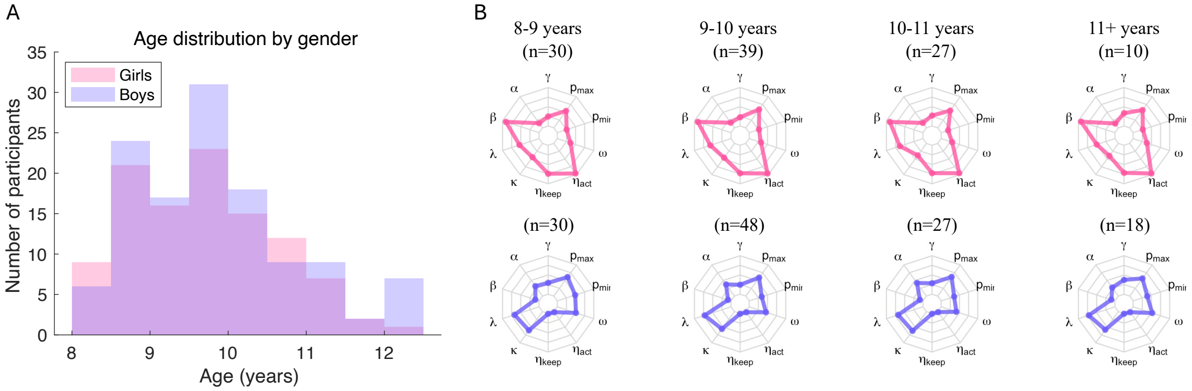


**Supplementary Figure 5. Age distribution and model parameter profiles by age and gender.** (A) Age distribution of participants by gender. Pink and blue histograms represent girls and boys, respectively. (B) Radar plots showing normalized model parameters for girls (top row) and boys (bottom row) across four age groups: 8–9 years ($8 \leq$ age $< 9$), 9–10 years ($9 \leq$ age $< 10$), 10–11 years ($10 \leq$ age $< 11$), and 11+ years (age $\geq 11$). Each axis corresponds to one parameter ($\gamma$, $\alpha$, $\beta$, $\lambda$, $\kappa$, $\eta_{\text{keep}}$, $\eta_{\text{act}}$, $\omega$, $p_{\min}$, $p_{\max}$). Parameter values were normalized across all participants using winsorized min–max scaling (5th–95th percentiles). Numbers above each panel indicate the sample size.

### Section 4: Supplementary Tables

**Table S1. Generalized linear mixed model results for behavioral decisions**

| Variable | Estimate | SE | $t$ | df | $p$ |
|---|---|---|---|---|---|
| (Intercept) | -1.96 | 0.24 | -8.10 | 22,868 | $5.8 \times 10^{-16***}$ |
| Scenario (punish–help) | -0.05 | 0.19 | -0.29 | 22,868 | 0.773 |
| Cost | -0.47 | 0.07 | -6.82 | 22,868 | $9.3 \times 10^{-12***}$ |
| Ratio | 0.11 | 0.09 | 1.25 | 22,868 | 0.212 |
| Inequality | 0.41 | 0.08 | 5.08 | 22,868 | $3.8 \times 10^{-7***}$ |
| Gender (boy–girl) | 0.54 | 0.33 | 1.66 | 22,868 | 0.097 |
| Scenario × Cost | 0.03 | 0.06 | 0.53 | 22,868 | 0.597 |
| Scenario × Ratio | -0.02 | 0.06 | -0.36 | 22,868 | 0.719 |
| Cost × Ratio | -0.17 | 0.04 | -3.97 | 22,868 | $7.1 \times 10^{-5***}$ |
| Scenario × Inequality | 0.04 | 0.06 | 0.66 | 22,868 | 0.509 |
| Cost × Inequality | 0.06 | 0.04 | 1.33 | 22,868 | 0.185 |
| Ratio × Inequality | -0.04 | 0.04 | -0.82 | 22,868 | 0.413 |
| Scenario × Gender | 0.29 | 0.25 | 1.14 | 22,868 | 0.256 |
| Cost × Gender | 0.29 | 0.09 | 3.22 | 22,868 | 0.001** |
| Ratio × Gender | 0.11 | 0.12 | 0.93 | 22,868 | 0.350 |
| Inequality × Gender | -0.17 | 0.11 | -1.58 | 22,868 | 0.115 |
| Scenario × Cost × Ratio | 0.06 | 0.06 | 0.96 | 22,868 | 0.338 |
| Scenario × Cost × Inequality | -0.07 | 0.06 | -1.18 | 22,868 | 0.240 |
| Scenario × Ratio × Inequality | 0.02 | 0.06 | 0.39 | 22,868 | 0.699 |
| Cost × Ratio × Inequality | -0.04 | 0.04 | -1.02 | 22,868 | 0.307 |
| Scenario × Cost × Gender | -0.02 | 0.08 | -0.19 | 22,868 | 0.851 |
| Scenario × Ratio × Gender | 0.21 | 0.08 | 2.53 | 22,868 | 0.011* |
| Cost × Ratio × Gender | 0.12 | 0.06 | 2.12 | 22,868 | 0.034* |
| Scenario × Inequality × Gender | 0.01 | 0.08 | 0.17 | 22,868 | 0.864 |
| Cost × Inequality × Gender | -0.06 | 0.06 | -1.03 | 22,868 | 0.304 |
| Ratio × Inequality × Gender | -0.03 | 0.06 | -0.44 | 22,868 | 0.657 |
| Scenario × Cost × Ratio × Inequality | -0.02 | 0.06 | -0.33 | 22,868 | 0.742 |
| Scenario × Cost × Ratio × Gender | -0.06 | 0.08 | -0.80 | 22,868 | 0.423 |
| Scenario × Cost × Inequality × Gender | 0.11 | 0.08 | 1.35 | 22,868 | 0.178 |
| Scenario × Ratio × Inequality × Gender | 0.10 | 0.08 | 1.19 | 22,868 | 0.234 |
| Cost × Ratio × Inequality × Gender | 0.05 | 0.06 | 0.95 | 22,868 | 0.344 |
| Scenario × Cost × Ratio × Inequality × Gender | -0.08 | 0.08 | -1.00 | 22,868 | 0.319 |

Scenario is a categorical predictor with two levels (punish vs. help). Gender is a categorical predictor with two levels (boy vs. girl). Inequality was calculated as the difference between the violator and the victim. All continuous predictors (cost, ratio, and inequality) were standardized using $z$-scores prior to model estimation. $^{*}p < 0.05$, $^{**}p < 0.01$, $^{***}p < 0.001$.

**Table S2. Linear mixed-effects model results for decision time**

| Variable | Estimate | SE | $t$ | df | $p$ |
|---|---|---|---|---|---|
| (Intercept) | 1.84 | 0.12 | 15.22 | 22,868 | $4.7 \times 10^{-52***}$ |
| Scenario (punish–help) | 0.23 | 0.12 | 1.92 | 22,868 | 0.055 |
| Cost | 0.10 | 0.05 | 1.88 | 22,868 | 0.060 |
| Ratio | 0.07 | 0.05 | 1.24 | 22,868 | 0.214 |
| Inequality | 0.04 | 0.05 | 0.85 | 22,868 | 0.395 |
| Gender (boy–girl) | -0.13 | 0.17 | -0.75 | 22,868 | 0.456 |
| Scenario × Cost | -0.22 | 0.07 | -2.99 | 22,868 | 0.003** |
| Scenario × Ratio | -0.02 | 0.07 | -0.27 | 22,868 | 0.791 |
| Cost × Ratio | 0.05 | 0.05 | 0.96 | 22,868 | 0.337 |
| Scenario × Inequality | -0.05 | 0.07 | -0.69 | 22,868 | 0.490 |
| Cost × Inequality | -0.06 | 0.05 | -1.14 | 22,868 | 0.254 |
| Ratio × Inequality | 0.05 | 0.05 | 1.00 | 22,868 | 0.320 |
| Scenario × Gender | -0.12 | 0.17 | -0.72 | 22,868 | 0.474 |
| Cost × Gender | -0.15 | 0.07 | -2.01 | 22,868 | 0.044* |
| Ratio × Gender | -0.13 | 0.07 | -1.76 | 22,868 | 0.078 |
| Inequality × Gender | 0.08 | 0.07 | 1.08 | 22,868 | 0.280 |
| Scenario × Cost × Ratio | -0.01 | 0.07 | -0.09 | 22,868 | 0.929 |
| Scenario × Cost × Inequality | 0.01 | 0.07 | 0.12 | 22,868 | 0.908 |
| Scenario × Ratio × Inequality | -0.08 | 0.07 | -1.05 | 22,868 | 0.294 |
| Cost × Ratio × Inequality | 0.07 | 0.05 | 1.35 | 22,868 | 0.177 |
| Scenario × Cost × Gender | 0.17 | 0.10 | 1.76 | 22,868 | 0.078 |
| Scenario × Ratio × Gender | 0.17 | 0.10 | 1.76 | 22,868 | 0.079 |
| Cost × Ratio × Gender | 0.00 | 0.07 | 0.06 | 22,868 | 0.955 |
| Scenario × Inequality × Gender | 0.00 | 0.10 | 0.03 | 22,868 | 0.975 |
| Cost × Inequality × Gender | 0.15 | 0.07 | 2.09 | 22,868 | 0.037* |
| Ratio × Inequality × Gender | -0.05 | 0.07 | -0.70 | 22,868 | 0.482 |
| Scenario × Cost × Ratio × Inequality | -0.02 | 0.07 | -0.30 | 22,868 | 0.767 |
| Scenario × Cost × Ratio × Gender | -0.07 | 0.10 | -0.72 | 22,868 | 0.472 |
| Scenario × Cost × Inequality × Gender | -0.14 | 0.10 | -1.40 | 22,868 | 0.163 |
| Scenario × Ratio × Inequality × Gender | 0.07 | 0.10 | 0.70 | 22,868 | 0.483 |
| Cost × Ratio × Inequality × Gender | -0.04 | 0.07 | -0.54 | 22,868 | 0.591 |
| Scenario × Cost × Ratio × Inequality × Gender | 0.04 | 0.10 | 0.39 | 22,868 | 0.696 |

Scenario is a categorical predictor with two levels (punish vs. help). Gender is a categorical predictor with two levels (boy vs. girl). Inequality was calculated as the difference between the violator and the victim. All continuous predictors (cost, ratio, and inequality) were standardized using $z$-scores prior to model estimation. $^{*}p < 0.05$, $^{**}p < 0.01$, $^{***}p < 0.001$.

**Table S3. Model comparison across all participants and by gender**

| Model | $ELPD_{LOO}$ | LOOIC |
|---|---|---|
| **All participants** | | |
| Baseline | -21310.83 | 42621.66 |
| SI | -22219.02 | 44438.03 |
| SI + SCI | -21265.55 | 42531.10 |
| SI + SCI + VCI | -20765.71 | 41531.42 |
| SI + SCI + VCI + EC | -20229.08 | 40458.16 |
| SI + SCI + VCI + EC + RP | -18194.81 | 36389.62 |
| SI + SCI + VCI + EC + RP + ID | -18123.49 | 36246.98 |
| **SI + SCI + VCI + EC + RP + ID + lapse** | **-17285.61** | **34571.23** |
| Heuristic | -17973.70 | 35947.40 |
| Heuristic + lapse | -17539.89 | 35079.78 |
| **Boys** | | |
| Baseline | -12014.99 | 24029.98 |
| SI | -12830.78 | 25661.57 |
| SI + SCI | -12548.63 | 25097.26 |
| SI + SCI + VCI | -12113.97 | 24227.93 |
| SI + SCI + VCI + EC | -11755.71 | 23511.43 |
| SI + SCI + VCI + EC + RP | -10400.52 | 20801.03 |
| SI + SCI + VCI + EC + RP + ID | -10376.97 | 20753.94 |
| **SI + SCI + VCI + EC + RP + ID + lapse** | **-9811.96** | **19623.92** |
| Heuristic | -10331.52 | 20663.04 |
| Heuristic + lapse | -9995.93 | 19991.86 |
| **Girls** | | |
| Baseline | -9295.84 | 18591.68 |
| SI | -9388.23 | 18776.46 |
| SI + SCI | -8716.92 | 17433.84 |
| SI + SCI + VCI | -8651.75 | 17303.49 |
| SI + SCI + VCI + EC | -8473.37 | 16946.74 |
| SI + SCI + VCI + EC + RP | -7794.30 | 15588.59 |
| SI + SCI + VCI + EC + RP + ID | -7746.52 | 15493.04 |
| **SI + SCI + VCI + EC + RP + ID + lapse** | **-7473.66** | **14947.31** |
| Heuristic | -7642.18 | 15284.36 |
| Heuristic + lapse | -7543.96 | 15087.92 |

$ELPD_{LOO}$ denotes the expected log predictive density estimated using PSIS-LOO. LOOIC is computed as $-2\times$ $ELPD_{LOO}$, with lower values indicating better predictive performance. Results are reported for all participants and separately for boys and girls.

**Table S4. Posterior estimates of gender differences (boys–girls) in group-level parameters**

| Parameter | Mean difference | 95% CrI | $p$ |
|---|---|---|---|
| $\alpha$ | 0.38 | [0.15, 0.62] | <0.001*** |
| $\eta_{\mathrm{act}}$ | -2.43 | [-6.91, 1.07] | 0.228 |
| $\eta_{\mathrm{keep}}$ | -2.04 | [-4.72, 0.08] | 0.073 |
| $\gamma$ | 0.10 | [-0.28, 0.46] | 0.555 |
| $\beta$ | -0.00 | [-0.02, 0.02] | 0.799 |
| $\kappa$ | 1.29 | [0.52, 2.12] | <0.001*** |
| $\lambda$ | 0.87 | [-4.53, 6.30] | 0.725 |
| $p_{\max}$ | 0.01 | [-0.04, 0.06] | 0.610 |
| $p_{\min}$ | 0.05 | [0.01, 0.08] | 0.016* |
| $\omega$ | 0.14 | [-0.02, 0.33] | 0.103 |

Mean difference indicates the posterior mean of the gender difference (boys–girls). CrI denotes the 95% Bayesian credible interval. $p$ represents the posterior probability that the parameter value is larger in boys than in girls. $^{*}p < 0.05$, $^{**}p < 0.01$, $^{***}p < 0.001$.

**Table S5. Statistical tests of cluster-level parameter differences for boys**

| Parameter | Test | Median [IQR] (C1) | Median [IQR] (C2) | $W$ | $r$ | $p_{\mathrm{corr}}$ |
|---|---|---|---|---|---|---|
| $\gamma$ | Wilcoxon | 1.819 [1.121] | 0.559 [0.874] | 2729 | 0.566 | $< 1.000 \times 10^{-6}$*** |
| $\alpha$ | Wilcoxon | 0.456 [0.485] | 0.770 [0.338] | 1029 | 0.279 | 0.012** |
| $\beta$ | Wilcoxon | 0.009 [0.000] | 0.009 [0.000] | 1698 | 0.053 | 1.000 |
| $\lambda$ | Wilcoxon | 6.385 [1.261] | 6.333 [1.423] | 1649 | 0.029 | 1.000 |
| $\kappa$ | Wilcoxon | 1.766 [1.387] | $-0.972$ [1.750] | 2716 | 0.559 | $< 1.000 \times 10^{-6}$*** |
| $\eta_{\mathrm{keep}}$ | Wilcoxon | 0.638 [0.007] | 0.640 [0.003] | 1271 | 0.159 | 0.390 |
| $\eta_{\mathrm{act}}$ | Wilcoxon | 1.208 [0.022] | 1.207 [0.011] | 1577 | 0.007 | 1.000 |
| $\omega$ | Wilcoxon | 0.534 [0.233] | 0.512 [0.173] | 1673 | 0.041 | 1.000 |
| $p_{\min}$ | Wilcoxon | 0.177 [0.079] | 0.076 [0.143] | 2382 | 0.393 | $1.040 \times 10^{-4}$*** |
| $p_{\max}$ | Wilcoxon | 0.193 [0.258] | 0.312 [0.054] | 835 | 0.376 | $2.170 \times 10^{-4}$*** |

Values are reported as median and interquartile range (IQR) for Cluster 1 (C1) and Cluster 2 (C2). Group differences were assessed using Wilcoxon rank-sum tests. $W$ denotes the Wilcoxon test statistic, and $r$ indicates the effect size ($r = Z/\sqrt{N}$). $p_{\mathrm{corr}}$ represents p-values corrected using the Holm–Bonferroni method.

**Table S6. Statistical tests of cluster-level parameter differences for girls**

| Parameter | Test | $H$ | $df$ | $\eta_H^2$ | $p_{\mathrm{corr}}$ |
|---|---|---|---|---|---|
| $\gamma$ | Kruskal–Wallis | 68.181 | 2 | 0.643 | $< 1.00 \times 10^{-6}$*** |
| $\alpha$ | Kruskal–Wallis | 27.460 | 2 | 0.247 | $1.10 \times 10^{-5}$*** |
| $\beta$ | Kruskal–Wallis | 7.696 | 2 | 0.055 | 0.213 |
| $\lambda$ | Kruskal–Wallis | 8.775 | 2 | 0.066 | 0.124 |
| $\kappa$ | Kruskal–Wallis | 26.293 | 2 | 0.236 | $2.00 \times 10^{-5}$*** |
| $\eta_{\mathrm{keep}}$ | Kruskal–Wallis | 20.465 | 2 | 0.179 | $3.60 \times 10^{-4}$*** |
| $\eta_{\mathrm{act}}$ | Kruskal–Wallis | 5.899 | 2 | 0.038 | 0.524 |
| $\omega$ | Kruskal–Wallis | 13.207 | 2 | 0.109 | 0.014* |
| $p_{\min}$ | Kruskal–Wallis | 32.480 | 2 | 0.296 | $1.00 \times 10^{-6}$*** |
| $p_{\max}$ | Kruskal–Wallis | 8.580 | 2 | 0.064 | 0.137 |

Kruskal–Wallis tests were used to assess overall cluster-level differences among girls. $H$ denotes the Kruskal–Wallis test statistic and $df$ indicates degrees of freedom. $\eta_H^2$ represents the nonparametric effect size for Kruskal–Wallis tests. $p_{\mathrm{corr}}$ represents p-values corrected using the Holm–Bonferroni method.

**Table S7. Post-hoc Dunn tests for cluster-level parameter differences among girls**

| Parameter | Comparison | $Z$ | $r$ | $p_{\mathrm{corr}}$ |
|---|---|---|---|---|
| $\gamma$ | Cluster 1 vs. Cluster 2 | -1.727 | -0.168 | 0.084 |
| $\gamma$ | Cluster 1 vs. Cluster 3 | 4.075 | 0.396 | $9.20 \times 10^{-5}$*** |
| $\gamma$ | Cluster 2 vs. Cluster 3 | 7.885 | 0.766 | $< 1.00 \times 10^{-6}$*** |
| $\alpha$ | Cluster 1 vs. Cluster 2 | -1.359 | -0.132 | 0.174 |
| $\alpha$ | Cluster 1 vs. Cluster 3 | -4.379 | -0.425 | $3.60 \times 10^{-5}$*** |
| $\alpha$ | Cluster 2 vs. Cluster 3 | -3.746 | -0.364 | $3.60 \times 10^{-4}$*** |
| $\kappa$ | Cluster 1 vs. Cluster 2 | 5.038 | 0.489 | $1.00 \times 10^{-6}$*** |
| $\kappa$ | Cluster 1 vs. Cluster 3 | 4.398 | 0.427 | $2.20 \times 10^{-5}$*** |
| $\kappa$ | Cluster 2 vs. Cluster 3 | -1.638 | -0.159 | 0.101 |
| $\eta_{\mathrm{keep}}$ | Cluster 1 vs. Cluster 2 | 1.341 | 0.130 | 0.180 |
| $\eta_{\mathrm{keep}}$ | Cluster 1 vs. Cluster 3 | -1.870 | -0.182 | 0.123 |
| $\eta_{\mathrm{keep}}$ | Cluster 2 vs. Cluster 3 | -4.423 | -0.430 | $2.90 \times 10^{-5}$*** |
| $\omega$ | Cluster 1 vs. Cluster 2 | 2.055 | 0.200 | 0.080 |
| $\omega$ | Cluster 1 vs. Cluster 3 | 3.528 | 0.343 | 0.001*** |
| $\omega$ | Cluster 2 vs. Cluster 3 | 1.606 | 0.156 | 0.108 |
| $p_{\min}$ | Cluster 1 vs. Cluster 2 | 1.068 | 0.104 | 0.285 |
| $p_{\min}$ | Cluster 1 vs. Cluster 3 | 4.518 | 0.439 | $1.90 \times 10^{-5}$*** |
| $p_{\min}$ | Cluster 2 vs. Cluster 3 | 4.356 | 0.423 | $2.60 \times 10^{-5}$*** |

Post-hoc pairwise comparisons were conducted using Dunn's tests following significant Kruskal–Wallis effects. Effect sizes are reported as $r = Z/\sqrt{N}$. $p_{\mathrm{corr}}$ $p_{\mathrm{corr}}$ represents p-values corrected using the Holm–Bonferroni method.

**Table S8. Generalized linear mixed model results including age effects**

| Variable | Estimate | SE | $t$ | df | $p$ |
|---|---|---|---|---|---|
| (Intercept) | -2.00 | 0.24 | -8.27 | 22,866 | $1.4 \times 10^{-16***}$ |
| Scenario (punish–help) | -0.06 | 0.19 | -0.32 | 22,866 | 0.753 |
| Cost | -0.48 | 0.07 | -6.97 | 22,866 | $3.3 \times 10^{-12***}$ |
| Ratio | 0.12 | 0.09 | 1.29 | 22,866 | 0.196 |
| Inequality | 0.40 | 0.08 | 5.04 | 22,866 | $4.8 \times 10^{-7***}$ |
| Age | -0.22 | 0.21 | -1.07 | 22,866 | 0.284 |
| Gender (boy–girl) | 0.55 | 0.33 | 1.68 | 22,866 | 0.092 |
| Scenario × Cost | 0.03 | 0.06 | 0.52 | 22,866 | 0.601 |
| Scenario × Ratio | -0.02 | 0.06 | -0.36 | 22,866 | 0.719 |
| Cost × Ratio | -0.17 | 0.04 | -3.95 | 22,866 | $7.8 \times 10^{-5***}$ |
| Scenario × Inequality | 0.04 | 0.06 | 0.67 | 22,866 | 0.503 |
| Cost × Inequality | 0.06 | 0.04 | 1.33 | 22,866 | 0.184 |
| Ratio × Inequality | -0.04 | 0.04 | -0.81 | 22,866 | 0.416 |
| Scenario × Gender | 0.29 | 0.25 | 1.16 | 22,866 | 0.244 |
| Cost × Gender | 0.31 | 0.09 | 3.37 | 22,866 | 0.001** |
| Ratio × Gender | 0.11 | 0.12 | 0.90 | 22,866 | 0.369 |
| Inequality × Gender | -0.16 | 0.11 | -1.50 | 22,866 | 0.134 |
| Age × Gender | 0.27 | 0.26 | 1.03 | 22,866 | 0.301 |
| Scenario × Cost × Ratio | 0.06 | 0.06 | 0.95 | 22,866 | 0.341 |
| Scenario × Cost × Inequality | -0.07 | 0.06 | -1.17 | 22,866 | 0.240 |
| Scenario × Ratio × Inequality | 0.02 | 0.06 | 0.38 | 22,866 | 0.703 |
| Cost × Ratio × Inequality | -0.04 | 0.04 | -1.02 | 22,866 | 0.307 |
| Scenario × Cost × Gender | -0.01 | 0.08 | -0.17 | 22,866 | 0.868 |
| Scenario × Ratio × Gender | 0.21 | 0.08 | 2.53 | 22,866 | 0.012* |
| Cost × Ratio × Gender | 0.12 | 0.06 | 2.10 | 22,866 | 0.036* |
| Scenario × Inequality × Gender | 0.01 | 0.08 | 0.18 | 22,866 | 0.860 |
| Cost × Inequality × Gender | -0.06 | 0.06 | -1.03 | 22,866 | 0.304 |
| Ratio × Inequality × Gender | -0.03 | 0.06 | -0.45 | 22,866 | 0.652 |
| Scenario × Cost × Ratio × Inequality | -0.02 | 0.06 | -0.33 | 22,866 | 0.742 |
| Scenario × Cost × Ratio × Gender | -0.06 | 0.08 | -0.80 | 22,866 | 0.424 |
| Scenario × Cost × Inequality × Gender | 0.11 | 0.08 | 1.34 | 22,866 | 0.180 |
| Scenario × Ratio × Inequality × Gender | 0.09 | 0.08 | 1.19 | 22,866 | 0.233 |
| Cost × Ratio × Inequality × Gender | 0.05 | 0.06 | 0.95 | 22,866 | 0.343 |
| Scenario × Cost × Ratio × Inequality × Gender | -0.08 | 0.08 | -1.00 | 22,866 | 0.318 |

Scenario is a categorical predictor with two levels (punish vs. help). Gender is a categorical predictor with two levels (boy vs. girl). Inequality was calculated as the difference between the violator and the victim. All continuous predictors (cost, ratio, inequality, and age) were standardized using $z$-scores prior to model estimation. $^{*}p < 0.05$, $^{**}p < 0.01$, $^{***}p < 0.001$.
